# Local cortical inhibitory subnetworks are shaped by pyramidal neuron progenitor type

**DOI:** 10.1101/2024.10.07.617021

**Authors:** Gemma Gothard, Kashif Mahfooz, Sarah E. Newey, Sasha Tinelli, Matthew J. Buchan, Richard J. Burman, Colin J. Akerman

## Abstract

The degree to which cortical neurons share inhibitory synaptic input determines their co-activity within a network. However, the principles by which inhibition is shared between neurons are not known. Here we combine *in utero* labeling with *in vivo* two-photon targeted patch-clamp recordings in mature cortex to reveal that a layer 2/3 (L2/3) pyramidal neuron’s local inhibitory input reflects the embryonic progenitor type from which the neuron is born. In contrast to neighboring neurons, pyramidal neurons derived from intermediate progenitors receive synaptic inhibition that is weakly coupled to local network activity. The underlying mechanisms do not depend upon the amount of inhibitory input received from different interneuron subclasses. Rather, progenitor type defines how much inhibitory input a neuron shares with its neighbors, which is reflected in how individual interneurons target pyramidal neurons according to progenitor type. These findings reveal new significance for progenitor diversity and identify ontogenetic origins of fine-scale inhibitory cortical subnetworks.

## Introduction

Synaptic circuits in the mammalian cerebral cortex are comprised from a heterogeneous array of excitatory glutamatergic pyramidal neurons and inhibitory GABAergic interneurons. While the exact wiring principles that generate functional circuits are not yet understood, the connections between excitatory and inhibitory neurons are critical for controlling the timing of action potentials, determining synaptic plasticity, and shaping population structure ^1–4^. Therefore, uncovering the fundamental wiring principles between excitatory and inhibitory neurons remains central to our understanding of cortical circuits.

Interneurons protect cortical networks from runaway excitation by interacting with excitatory neurons through particular circuit motifs ^5^. Early evidence described a circuit motif referred to as ‘blanket inhibition’, whereby inhibitory interneurons connect to all surrounding excitatory neurons without specificity ^6,7^. A series of more recent studies, however, support the idea that cortical pyramidal neurons receive selective input from interneurons ^8–15^. This selectivity in synaptic inhibition is thought to generate ‘subnetworks’ of functionally-related neurons within cortex, which can be defined by their local and long-range patterns of synaptic connectivity ^8,9,11–13^. These subnetworks serve to separate information processing streams within cortical circuits, such that neurons in one subnetwork can more easily decorrelate their activity from neurons in a separate subnetwork ^3,11^. This is advantageous as it increases the computational capacity of cortex and can generate multiple processing streams for different types of information ^8–13,16^. Despite the functional significance of cortical subnetworks, the processes by which these are established are not known.

The connectivity of mature cortical pyramidal neurons is reflected in their developmental trajectories in the embryonic brain ^17–22^. During neurogenesis in the mouse, excitatory neurons are born from a pool of progenitor cells within the ventricular and subventricular zones of the dorsal telencephalon ^23–25^. The principal neurogenic cell type is the radial glial cell (RGC) which firstly undergo symmetric divisions, producing two daughter RGCs that amplifies the RGC pool ^26^. At around embryonic day 12 (E12) RGCs switch to self-renewing asymmetric divisions, producing one RGC and either a post-mitotic daughter neuron or an intermediate progenitor (IP) ^26,27^. IPs then terminally divide once to produce two post-mitotic daughter neurons ^24,28^.The time at which L2/3 neurons are born, pyramidal neurons can either be born from self-renewing divisions by RGCs, or from terminal divisions by IPs ^29–34^. These distinct trajectories have implications for shaping the excitatory synaptic connectivity of mature cortical neurons ^34–38^. However, it is unknown whether progenitor type influences the inhibitory connections that a L2/3 pyramidal neuron receives and therefore shapes the formation of inhibitory subnetworks within cortex.

Here we demonstrate that a L2/3 pyramidal neuron’s inhibitory subnetwork depends upon the embryonic progenitor type from which the pyramidal neuron is derived. Employing *in utero* fate mapping techniques to label cohorts of pyramidal neurons, we then use *in vivo* two-photon targeted whole-cell patch-clamp recordings to reveal that progenitor type predicts how a pyramidal neuron’s synaptic inhibition relates to local network activity. This is not associated with differences in the level of inhibition that the neuron receives from interneuron subclasses. Instead, we show that progenitor type defines the degree to which pyramidal neurons receive input from the same interneurons as their neighbors, with PV interneurons differentially targeting pyramidal neurons based on the post-synaptic neuron’s progenitor type.

## Results

### Progenitor type defines how strongly coupled a L2/3 pyramidal neuron’s synaptic inhibition is to local network activity

Mouse cortical excitatory neurons and inhibitory interneurons are born from spatially separate pools of embryonic progenitors in the pallium and subpallium, respectively (**Fig. 1a**). These neurons then migrate to meet in the cortical plate, where they form synaptic circuits over the course of development ^39–43^ (**Fig. 1b**). The pallial progenitor cells that generate excitatory neurons are heterogenous, such that neighboring L2/3 pyramidal neurons can be generated directly from RGCs or indirectly from IPs ^24,44–47^. To target pallial progenitor cells giving rise to L2/3 pyramidal neurons, we performed *in utero* electroporation (IUE) in C57BL/6 mice at embryonic day 15.5 (E15.5) (**Fig. 1c**) ^48^. Consistent with previous work, the tubulin alpha1 (Tα1) promoter was used to label a subset of L2/3 pyramidal neurons derived from a population of intermediate progenitors (IPs) ^34,38,49–51^, which was shown to include both apical and basal IPs (**Supplementary Fig. 1**). Our labeling strategy used Cre recombinase downstream of the Tα1 promoter to permanently turn on expression of enhanced green fluorescent protein (GFP) from a reporter construct (**Fig. 1c**). The region corresponding to primary somatosensory cortex (S1) was targeted, such that 24 hours after IUE, GFP-expressing IPs and newly born neurons derived from IPs (‘IP-derived’) were observed in the germinal zone (**Fig. 1d**). When the mice developed to 5 weeks of age, GFP-expressing IP-derived pyramidal neurons were observed in L2/3 of mature S1 (**Fig. 1d**; average depth from pial surface: 233.28 ± 8.43 µm).

To investigate whether progenitor type influences a L2/3 pyramidal neuron’s synaptic inhibition, we conducted *in vivo* two-photon targeted whole-cell patch-clamp recordings of inhibitory synaptic conductances from the IP-derived neurons (**Fig. 1e**; see Methods). Patch-clamp recordings were combined with simultaneous local field potential (LFP) recordings to be able to relate the neuron’s synaptic inhibition to local network activity (**Fig. 1e; Supplementary Fig. 2**) ^52^. GFP-expressing IP-derived L2/3 neurons were targeted for whole-cell patch-clamp recordings and compared to recordings from unlabeled L2/3 pyramidal neurons, which were therefore derived from a mix of different lineages. In a subset of recordings, neurons were filled with biocytin and visualised post-hoc to confirm their pyramidal identity (**Fig. 1f**).

Consistent with previous evidence that L2/3 pyramidal neurons are under strong inhibitory synaptic control *in vivo*, our whole-cell patch-clamp recordings revealed robust inhibitory synaptic conductances in all of the recorded neurons (n = 22 neurons in 10 animals) ^52^. We observed that inhibitory synaptic inputs in unlabeled neurons were typically associated with rapid negative deflections in the LFP, consistent with periods of local network activity ^52–54^. However, inhibitory synaptic inputs in IP-derived neurons were often associated with less activity in the LFP (**Fig. 1g**). This suggests that the inhibitory synaptic input to IP-derived neurons is weakly correlated with local network activity. To capture this relationship, we applied a measure of population coupling that has been used to relate an individual neuron’s activity to the activity of the local network measured from the LFP ^53,54^. For each neuron, the onset times of every inhibitory synaptic input was identified via automated detection and used to extract the surrounding 100 ms LFP window. By averaging these LFP windows, it was possible to define an input-triggered LFP (IT-LFP; **Fig. 1h**), which reflects the average local network activity during any given inhibitory synaptic input that a neuron receives (**Fig. 1h**). The peak IT-LFP (defined as the mean of the IT-LFP during the 10 ms following the onset of the inhibitory input) was significantly smaller in IP-derived neurons than unlabeled neurons, revealing that synaptic inhibition in IP-derived neurons was more weakly coupled to local network activity (**Fig. 1h**; unlabeled IT-LFP: -245.35 ± 30.62 µV, IP-derived IT-LFP: -170.09 ± 16.27 µV). This observation was not associated with any differences in the distribution of LFP values during recordings from IP-derived and unlabeled neurons (**Supplementary Fig. 3**). Nor were there differences in the frequency or amplitude of inhibitory synaptic conductances between the neuronal populations (**Supplementary Fig. 4**). These data suggest that L2/3 pyramidal neurons receive similar overall levels of inhibitory synaptic input, but that the inhibition received by an IP-derived L2/3 pyramidal neuron is more weakly coupled to local network activity.

**Figure 1:**
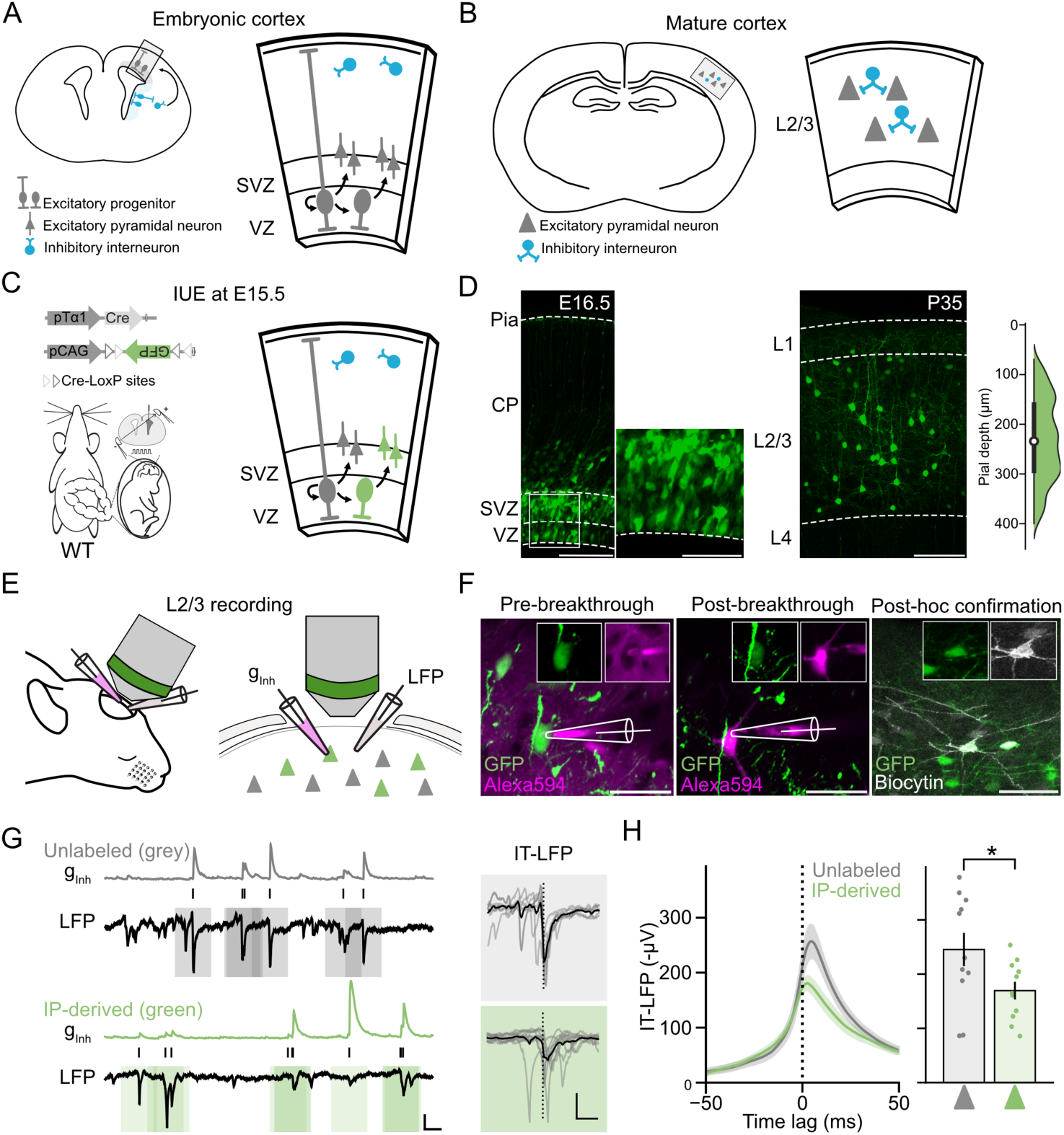
Progenitor type defines how strongly coupled a L2/3 pyramidal neuron’s synaptic inhibition is to local network activity *in vivo*. (**a**) Excitatory pyramidal neurons and inhibitory interneurons born from progenitor pools in the pallium and subpallium, respectively, migrate to meet in the cortical plate (CP). VZ, ventricular zone, SVZ, subventricular zone. (**b**) The excitatory neurons and inhibitory interneurons go on to generate subnetworks within mature cortex. (**c**) IUE of pTα1-Cre and pCAG-FLEX-GFP was used to label IP-derived L2/3 pyramidal neurons in S1. (**d**) GFP-expressing IPs and newly born IP-derived neurons were observed 24 h after IUE (left). Scale bar, 100 µm or 50 µm. At P35, IP-derived pyramidal neurons were distributed throughout L2/3 (mean pial depth: 233.28 ± 0.85 µm, n = 397 neurons from 4 animals) (right). Scale bar, 100 µm. (**e**) At P28-42, animals underwent *in vivo* electrophysiology that combined simultaneous whole-cell patch-clamp recordings of inhibitory synaptic conductances (gInh) and the local field potential (LFP). (**f**) Example two-photon targeted whole-cell patch-clamp recording from a GFP-expressing IP-derived L2/3 pyramidal neuron with post-hoc confirmation. Scale bar, 50 µm. (**g**) Example simultaneous recordings of inhibitory synaptic inputs and the LFP for an unlabeled (top) and IP-derived (bottom) L2/3 pyramidal neuron. The onset of individual inhibitory inputs were identified via automated detection (vertical ticks) and the surrounding 500 ms of LFP was extracted (shaded areas), such that an input-triggered LFP (IT-LFP) was generated for each input (right). Scale bar, 2 nS, 200 µV, 250 ms. IT-LFP inset scale bar, 200 µV and 100 ms. (**h**) IT-LFP averaged across all neurons within each group for the 100 ms surrounding inhibitory input onset (left). IP-derived neurons exhibited a lower peak IT-LFP than unlabeled neurons (right) (two-tailed unpaired t-test, *n* = 11 unlabeled neurons from 5 animals, *n* = 11 IP-derived neurons from 5 animals, *P* = 0.042, df = 20, T = 2.17).

To support these observations, we performed a separate event-based analysis in which we used automated methods to detect the local network events (defined as the peak of deflections identified in the LFP recordings) and the inhibitory synaptic events (defined as the onset of each synaptic input, **Fig. 2a**). This revealed no differences in frequency or amplitude of LFP network events or synaptic events between the two neuronal populations (**Supplementary Fig. 4**). To then capture the temporal relationship between local network events and inhibitory synaptic events, we calculated an event time tiling coefficient (ETTC), which measures the synchrony of local network events and synaptic events over short timescales (**Fig. 2a**; see Methods)^55^. For our analysis, we set this timescale as 25 ms, meaning the ETTC increases each time inhibitory synaptic events and LFP events occur within 25 ms of one another. We confirmed that ETTC was stable over the course of a single recording (**Fig. 2b**; first half of recording versus second half of recording: r^2^ = 0.94). When this analysis was used to compare the different neuronal populations, it was found that IP-derived neurons had a lower ETTC than unlabeled neurons (**Fig. 2c** and **2d**; unlabeled ETTC: 0.54 ± 0.05, IP-derived ETTC: 0.38 ± 0.05), revealing that local network events are less likely to co-occur with synaptic inhibition in IP-derived neurons. This supports the conclusion that progenitor type can define how strongly coupled a L2/3 pyramidal neuron’s synaptic inhibition is to local network activity.

**Figure 2:**
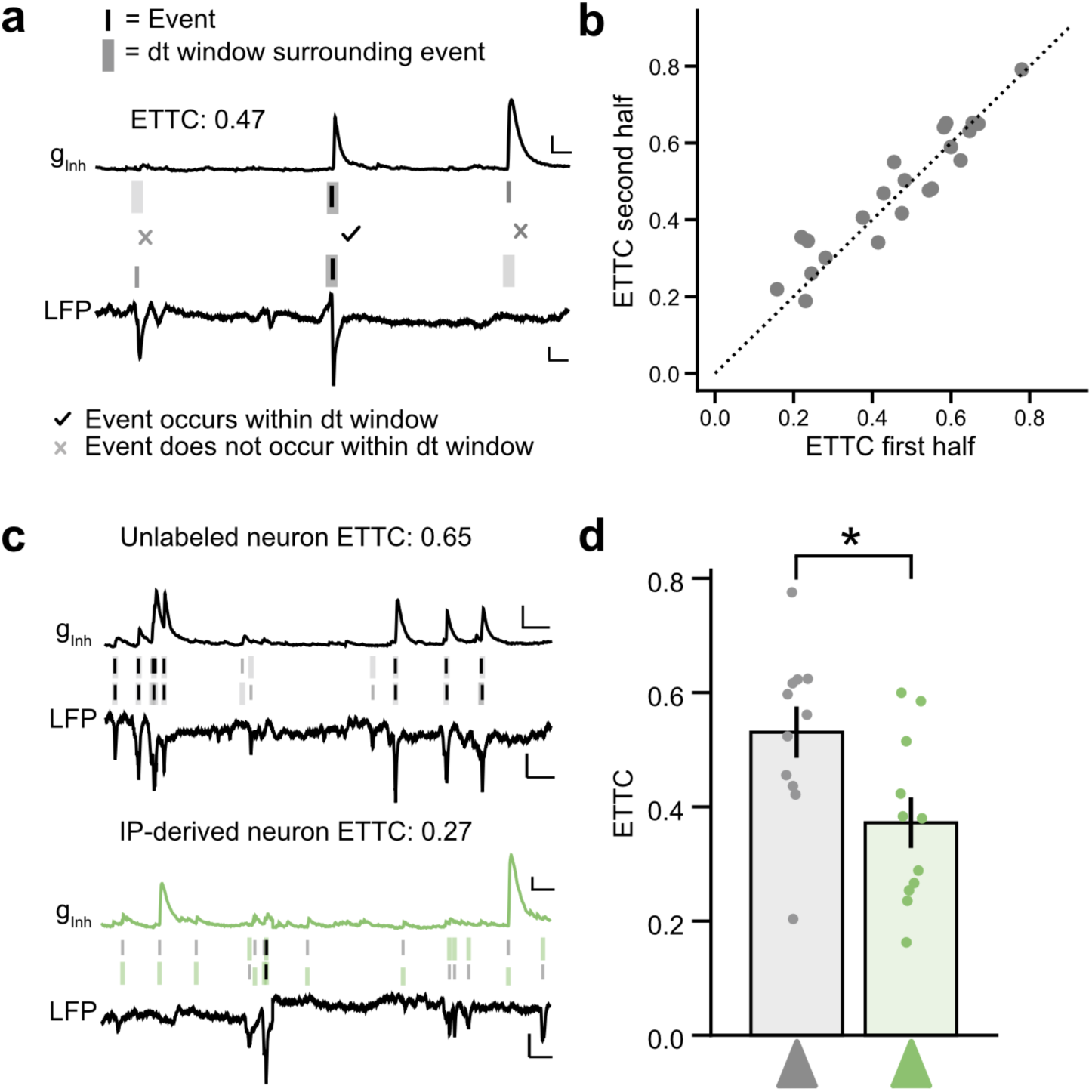
Progenitor type predicts the correlation between a L2/3 pyramidal neuron’s inhibitory synaptic events and local network events *in vivo*. (**a**) Example traces depicting detection of inhibitory synaptic events (top) and local network events (bottom). A 25 ms ± dt window was used to determine the event time tiling coefficient (ETTC) - a bidirectional correlation measure of the synaptic events and local network events. ETTC increases when inhibitory synaptic events and LFP events occur close to one another in time (see example with tick) and decreases when inhibitory synaptic events and LFP events occur independently of one another (see examples with crosses). Vertical lines indicate a detected event in the synaptic recording or the LFP. Shaded areas indicate the ± 25 ms dt window that was used to search for an event in the other trace. Darker vertical lines and shaded areas indicate where inhibitory events and network events co-occur within ± 25 ms of one another. Scale bar, 1 nS, 100 µV, 125 ms. (**b**) ETTC values for individual neurons were stable between the first and second half of the recording (*n* = 22 neurons from 10 animals, r^2^ = 0.94). (**c**) Example trace for an unlabeled (top, grey) and an IP-derived (bottom, green) L2/3 pyramidal neuron with their respective ETTC values. Scale bar, 1 nS, 100 µV, 250 ms. (**d**) ETTC was lower for IP-derived neurons than unlabeled neurons (two-tailed Mann-Whitney U test, *n* = 11 unlabeled neurons from 5 animals, *n* = 11 IP-derived neurons from 5 animals, *P* = 0.021, U = 97).

### Progenitor type does not determine a L2/3 pyramidal neuron’s level of overall input from different interneuron subclasses

To understand how progenitor type is able to define a L2/3 pyramidal neuron’s coupling to the local inhibitory network, we considered potential underlying cellular mechanisms. One explanation is that L2/3 pyramidal neurons from different progenitor types are specified to receive different amounts of input from distinct interneuron subclasses. To investigate this possibility, an optogenetic approach was used to determine whether the input from interneuron subclasses varies depending on the pyramidal neuron’s progenitor type. To be compatible with Cre-Lox mice that express Channelrhodopsin (ChR2) in defined interneuron populations, we developed a dual-color Flpo recombinase reporter system (pEf1a-FRT-tdTom/EGFP, see **Methods**) that distinguished two different subpopulations of L2/3 pyramidal neurons based on their progenitor type. Flpo recombinase downstream of the Tα1 promoter was combined with a reporter construct incorporating a flexible excision (FLEx) cassette, in which Flpo recombination permanently switches expression from TdTomato fluorescent protein to GFP (**Fig. 3a**). With this strategy, the IP-derived pyramidal neurons express GFP, whilst neighboring pyramidal neurons derived from other progenitors (OPs) express TdTomato (**Fig. 3b** and **3c**) ^56^. “OP-derived” was therefore used to indicate neurons that are derived from progenitors in which the Tα1 promoter was not active. The OP-derived L2/3 pyramidal neurons were shown to include neurons from a Glast-positive lineage, consistent with this population including neurons derived directly from radial glial cells (**Supplementary Fig. 5**) ^34,45^. Also in line with previous work, the soma of IP-derived and OP-derived L2/3 pyramidal neurons showed overlapping distributions in mature cortex, with the average IP-derived neuron located slightly further from pia, consistent with the OP population including progenitors that undergo self-renewing divisions (**Fig. 3c**; OP-derived: 217.93 ± 6.74 µm, IP-derived: 257.56 ± 10.58 µm) ^31,56^.

Our dual-color reporter meant that simultaneous recordings could be performed from IP-derived and neighboring OP-derived L2/3 pyramidal neurons in order to quantify the relative amount of inhibitory input in the same tissue. To ensure differences in pial depth did not influence our results, our recordings targeted IP-derived and OP-derived neurons that were equidistant from the pia. Our first experiment examined PV interneurons, which are known to provide strong peri-somatic input to L2/3 pyramidal neurons (**Fig. 3d**) ^57–59^. Animals that had undergone IUE were allowed to reach maturity (ages P21-42) and acute slices were prepared from S1 to allow for simultaneous whole-cell patch-clamp recordings from neuronal pairs comprising a GFP-expressing IP-derived L2/3 pyramidal neuron and a neighboring TdTomato-expressing OP-derived L2/3 pyramidal neuron (mean inter-somatic distance of 67.38 ± 7.21 µm). Brief pulses from a widefield light-emitting diode (LED; 473 nm, 1 ms duration) were used to activate ChR2-expressing PV interneurons throughout the brain slice, whilst the resulting monosynaptic inhibitory post-synaptic currents (IPSC_PV_) were recorded and their amplitudes compared (**Fig. 3d-f**). Consistent with previous work, this elicited a large IPSC_PV_ whose amplitude varied across neurons ^59^. However, there was no systematic difference in the amplitude of the IPSC_PV_ between IP-derived and OP-derived neurons (**Fig. 3e**; OP-derived: 462.71 ± 59.32 pA, IP-derived: 444.00 ± 69.92 pA). This was similar to randomly sampled pairs of unlabeled neurons (**Supplementary Fig. 6**), suggesting that progenitor type is not associated with a different level of PV interneuron input.

**Figure 3:**
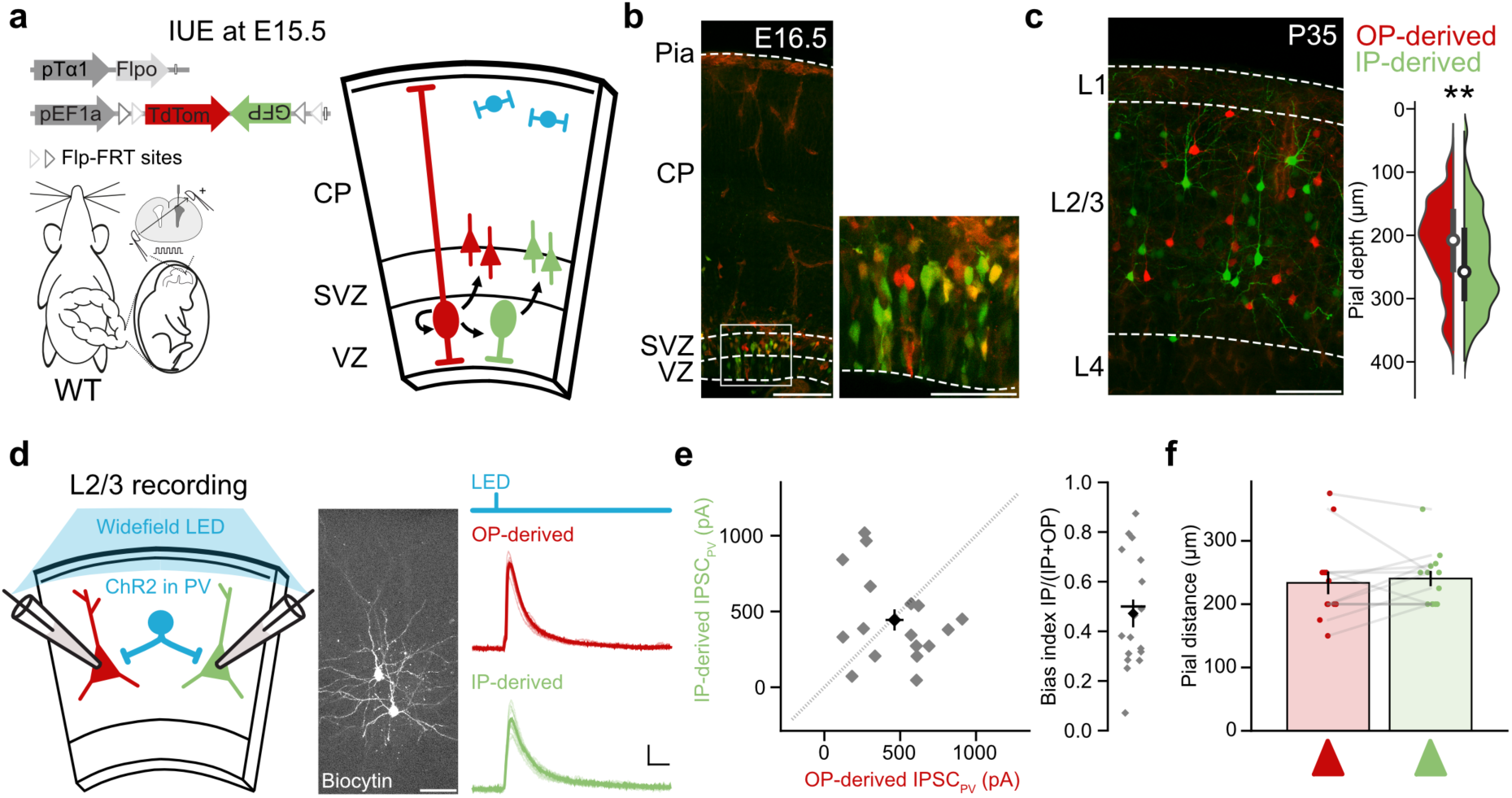
Progenitor type does not determine a L2/3 pyramidal neuron’s level of input from PV interneurons. (**a**) IUE at E15.5 was used to label IP-derived and OP-derived L2/3 pyramidal neurons in mice expressing ChR2 in PV interneurons. (**b**) TdTomato-expressing OPs and GFP-expressing IPs can be seen along with newly born IP-derived and OP-derived neurons 24 hours following IUE. Scale bar, 100 µm, 50 µm for inset. (**c**) At P35, the L2/3 pyramidal neurons showed overlapping distributions, with the average IP-derived neuron further from pia than the average OP-derived neuron (two-tailed paired t-test, *n* = 679 neurons from 14 animals, *P* = 0.01, df = 13, T = -3.02). Pial depth is shown as a kernel density estimate alongside mean ± standard deviation. Scale bar, 100 µm. (**d**) At P21, acute slices were prepared from S1 and simultaneous recordings were performed from pairs of IP-derived and OP-derived L2/3 pyramidal neurons, whilst brief widefield light pulses from a LED (473 nm, 1 ms duration) were used to activate ChR2-expressing PV interneurons throughout the brain slice (left). In a subset of recordings, neurons were filled with biocytin and visualised post-hoc to confirm pyramidal morphology (middle). Scale bar, 100 µm. Example IPSCPV recorded from an IP-derived and OP-derived neuron pair (right). Scale bar, 100 pA, 25 ms. (**e**) IPSCPV did not differ between IP-derived and OP-derived neurons (one sample t-test, *n* = 17 pairs from 8 animals, *P* = 0.80, df = 16, T = 0.26). (**f**) The recorded IP-derived and OP-derived neurons were at similar distances from pia (two-tailed Wilcoxon signed-rank test, OP-derived mean: 233.62 ± 17.98 µm, IP-derived mean: 240.46 ± 12.13 µm, *n* = 13 pairs from 7 animals, *P* = 0.43, df = 12, W = 51.5).

We next considered whether progenitor type predicts the amount of input from SST interneurons, which target the dendrites of L2/3 pyramidal neurons ^60^. Widefield ChR2 activation of SST interneurons elicited a large IPSC_SST_ in pairs of neighboring L2/3 pyramidal neurons (**Fig. 4a**; mean inter-somatic distance of 85.55 ± 10.12 µm), but the IPSC_SST_ amplitude did not exhibit systematic differences between IP-derived and OP-derived neurons (**Fig. 4b** and **4c**; OP-derived: 497.47 ± 42.12 pA, IP-derived: 474.43 ± 48.32 pA), similar to pairs of unlabeled neurons (**Supplementary Fig. 6**). Finally, we examined the input from VIP interneurons, which provide monosynaptic input to L2/3 pyramidal neurons, in addition to their role in disinhibitory cortical circuits ^61,62^. Widefield ChR2 activation of VIP interneurons tended to elicit somewhat smaller post-synaptic responses (**Fig. 4d**; mean inter-somatic distance of 67.25 ± 6.90 µm), but again the amplitude of IPSC_VIP_ did not differ between IP-derived and OP-derived neurons (**Fig. 4e** and **4f**; OP-derived: 182.19 ± 40.313, IP-derived: 153.20 ± 18.66 pA), as was the case for randomly sampled pairs of unlabeled neurons (**Supplementary Fig. 6**). These results indicate that the overall amount of synaptic inhibition provided by PV, SST or VIP interneurons does not vary according to a L2/3 pyramidal neuron’s progenitor type. Consequently, the progenitor-dependent differences in how a L2/3 pyramidal neuron couples to the local inhibitory network are unlikely to reflect different input levels from the major interneuron subclasses.

**Figure 4:**
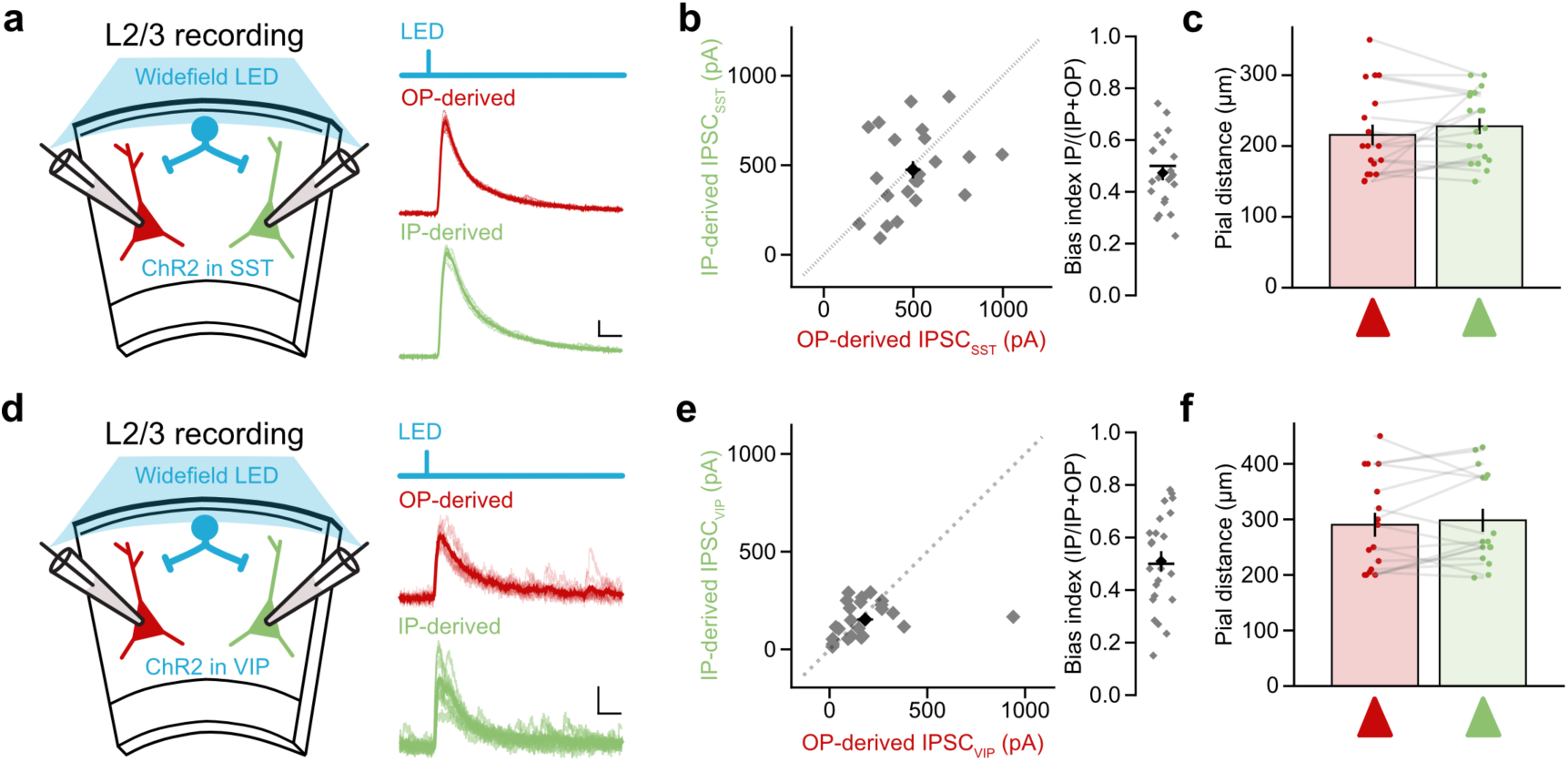
Progenitor type does not determine a L2/3 pyramidal neuron’s level of input from SST or VIP interneurons. (**a**) Simultaneous patch-clamp recordings were performed from IP-derived and OP-derived L2/3 pyramidal neurons, whilst brief widefield light pulses from a LED (473 nm, 1 ms duration) were used to activate ChR2-expressing SST interneurons (left). Example IPSCSST recorded from an IP-derived and OP-derived pair (right). Scale bar, 100 pA, 25 ms. (**b**) IPSCSST did not differ between IP-derived and OP-derived neurons (one-sample t-test, *n* = 22 pairs from 11 animals, *P* = 0.36, df = 21, T = -0.94). (**c**) Recorded IP-derived and OP-derived neurons were equidistant from pia (two-tailed Wilcoxon signed-rank test, OP-derived mean: 219.06 ± 14.56 µm, IP-derived mean: 226.72 ± 11.76 µm, *n* = 18 pairs from 11 animals, *P* = 0.48, W = 54.5). (**d**) Simultaneous patch-clamp recordings were made from IP-derived and OP-derived neurons, whilst activating ChR2-expressing VIP interneurons (left). Example IPSCVIP recorded from an IP-derived and OP-derived pair (right). Scale bar, 50 pA, 25 ms. (**e**) IPSCVIP amplitude did not differ between IP-derived and OP-derived neurons (one-sample t-test, *n* = 23 pairs from 7 animals, *P* = 0.80, df = 22, T = 0.26). (**f**) Recorded IP-derived and OP-derived neurons were at similar distances from pia (two-tailed Wilcoxon signed-rank test, OP-derived mean: 290.31 ± 21.63 µm, IP-derived mean: 298.44 ± 20.80 µm, n = 16 pairs from 7 animals, *P* = 0.43, W = 51.5).

### Progenitor type specifies a L2/3 pyramidal neuron’s local inhibitory subnetwork

We therefore considered an alternative cellular mechanism by which progenitor type could define how strongly a L2/3 pyramidal neuron’s synaptic inhibition couples to local network activity. We wondered whether progenitor type may specify how pyramidal neurons share input from the same individual presynaptic interneuron. To explore this, our first approach was to investigate the degree to which spontaneous inhibitory inputs are shared by neighboring L2/3 pyramidal neurons, as this can be used to measure common input from the same individual pre-synaptic interneuron ^63^. Following IUE at E15.5 with the dual-color reporter constructs, animals were allowed to reach maturity (ages P21-P42) and acute slices were prepared from S1. Simultaneous whole-cell patch-clamp recordings were performed from pairs of neurons comprising either two unlabeled neighboring pyramidal neurons, whose progenitor type was therefore unknown (**Fig. 5a-c**), or pairs comprising an IP-derived and a neighboring OP-derived pyramidal neuron (**Fig. 5d-f**). Spontaneous IPSCs were observed in all recorded neurons and an input threshold was used to maximize the detection of unitary, action potential-evoked IPSCs, rather than miniature events (see Methods) ^59,64^.

**Figure 5:**
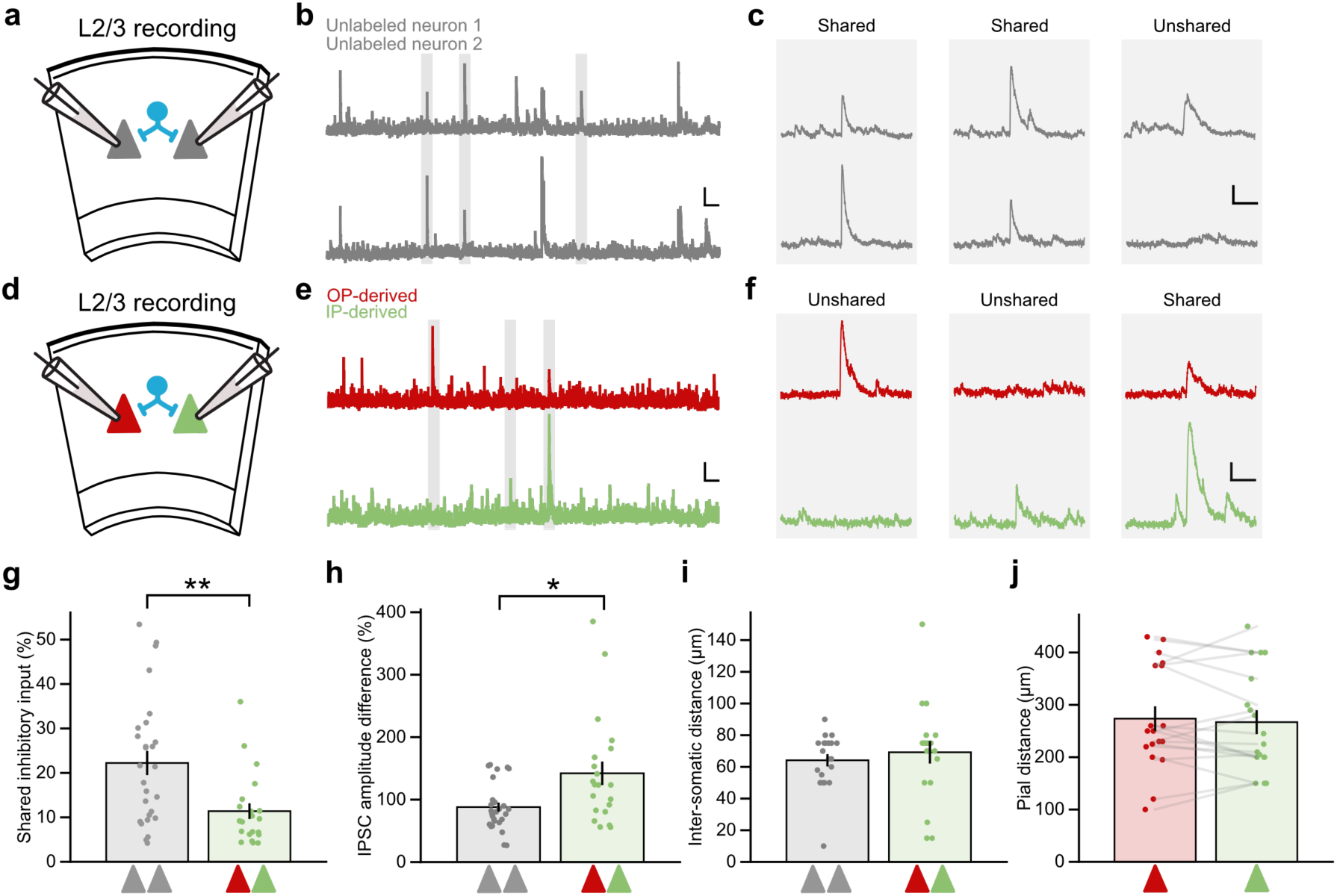
Progenitor type predicts how neighboring L2/3 pyramidal neurons share inhibitory synaptic input. (**a**) Simultaneous whole-cell patch-clamp recordings from pairs of neighboring unlabeled pyramidal neurons at P21-42 were used to detect spontaneous action-potential-evoked IPSCs. (**b**) Example recordings as described in ‘a’. Scale bar, 50 pA, 1 s. (**c**) Expanded shaded regions from ‘b’, illustrating examples of shared and unshared inhibitory inputs. Scale bar, 50 pA, 100 ms. (**d**) Simultaneous recordings from pairs of neurons comprising an IP-derived and a neighboring OP-derived pyramidal neuron. (**e**) Example recordings as described in ‘d’. Scale bar, 50 pA, 1 s. (**f**) Expanded shaded regions from ‘e’, illustrating examples of shared and unshared inhibitory inputs. Scale bar, 50 pA, 100 ms. (**g**) Percentage of shared inhibitory inputs was lower for pairs comprising an OP-derived and an IP-derived L2/3 pyramidal neuron, than for pairs comprising two unlabeled L2/3 pyramidal neurons (two-tailed Mann-Whitney U test, *n* = 27 pairs of unlabeled neurons from 16 animals, *n* = 21 pairs of IP-derived and OP-derived neurons from 10 animals, *P* = 0.004, U = 421). (**h**) IPSC amplitude difference for shared inhibitory inputs (expressed as a percentage of median shared IPSC amplitude) was greater for pairs comprising an IP-derived and OP-derived L2/3 pyramidal neuron, than for pairs comprising two unlabeled L2/3 pyramidal neurons (two-tailed Mann-Whitney U test, unlabeled pairs: 88.02 ± 7.28 %, *n* = 27 pairs from 16 animals, IP-derived and OP-derived pairs: 142.26 ± 18.85 %, *n* = 21 pairs from 10 animals, *P* = 0.01, U = 166). (**i**) Inter-somatic distance was not different between the pair types (two-tailed Mann-Whitney U test, unlabeled mean: 64.15 ± 3.90 µm, *n* = 20 pairs of unlabeled neurons from 13 animals, IP-derived and OP-derived mean: 69.21 ± 7.14 µm, *n* = 19 pairs of from 11 animals, *P* = 0.48, U = 164.5). (**j**) Distance from pia was not different between IP-derived and OP-derived neurons (two-tailed paired t-test, OP-derived neurons: 266.94 ± 22.80 µm, IP-derived neurons: 273.61 ± 23.65 µm, *n* = 18 pairs from 11 animals, *P* = 0.53, df = 17, T = 0.64).

The amplitude and frequency of IPSCs were comparable for the different neuronal populations (**Supplementary Fig. 7**). However, an analysis of the relative timing of each input allowed us to classify whether an input was shared (received simultaneously by both neurons) or unshared (received by only one neuron) between the recorded pair (**Fig. 5c** and **5f**). In pairs of unlabeled pyramidal neurons, 22.23 ± 2.73 % of inhibitory inputs were classified as shared. In contrast, shared inhibitory inputs were much less common in pairs comprising IP-derived and OP-derived pyramidal neurons, where the proportion of shared inputs was only 11.51 ± 1.81 % (**Fig. 5g**). Furthermore, whilst the mean amplitude of shared inhibitory inputs were similar (**Supplementary Fig. 8**), the amplitude of shared inputs was more variable for IP-derived and OP-derived pairs than for unlabeled pairs (**Fig. 5h**; unlabeled neuron pairs: 88.02 ± 7.28 %, IP-derived and OP-derived neuron pairs: 142.26 ± 18.85 %; where the difference in shared input amplitude is expressed as a percentage of the median shared input amplitude). Thus, pyramidal neurons from different progenitor types had fewer shared inhibitory inputs and the shared inputs they did have were more variable in strength. The inter-somatic distances of the neuronal pairs were comparable (**Fig. 5i**; unlabeled pairs: 64.15 ± 3.90 µm, IP-derived and OP-derived pairs: 69.21 ± 7.14 µm), as were the distances from pia for the IP-derived and OP-derived neurons (**Fig. 5j**; OP-derived neurons: 266.94 ± 22.80 µm, IP-derived neurons: 273.61 ± 23.65 µm; **Supplementary Fig. 8**). We also confirmed that the difference in the proportion of shared inputs was evident across a range of different input detection thresholds (**Supplementary Fig. 8**).

These data indicate that progenitor type can specify the degree to which neighboring pyramidal neurons share their inhibitory synaptic input from presynaptic interneurons. If this is the mechanism by which progenitor type defines a L2/3 pyramidal neuron’s coupling to the local inhibitory network, we hypothesised that one should be able to detect biased output in terms of how individual interneurons target L2/3 pyramidal neurons. To investigate this, we designed an optogenetic experiment to assess the output from putative single PV interneurons onto IP-derived and OP-derived neurons. Monosynaptic IPSCs were recorded from simultaneously recorded pairs of neighboring L2/3 pyramidal neurons, except now a laser beam was used to focally activate ChR2-expressing PV interneurons with spatially-restricted pulses of light (**Fig. 6a-c**; see Methods). In line with previous ChR2-assisted mapping of unitary connectivity ^65^, the photostimulation conditions were set such that a mixture of successes and failures in post-synaptic responses were observed across trials, consistent with the activation of a single or small number of interneurons (see Methods). If interneurons connect with similar probability and strength to the two pyramidal neurons, one would expect to observe similar post-synaptic responses on a trial-to-trial basis. In contrast, if interneurons exhibit a biased output, the expectation is that the post-synaptic responses would exhibit differences on a trial-to-trial basis (**Fig. 6a-c**). We first asked what proportion of trials exhibited responses in both neurons (i.e. a shared output) and found that the probability of shared output responses was lower in pairs comprising an IP-derived and a neighboring OP-derived pyramidal neuron, than pairs of unlabeled L2/3 pyramidal neurons (**Fig. 6d**; unlabeled pairs: 0.68 ± 0.07, IP-derived and OP-derived pairs: 0.48 ± 0.05). Furthermore, we observed that progenitor type was associated with greater variability in shared output responses. IP-derived and OP-derived pairs exhibited greater differences in response amplitudes than unlabeled pairs (**Fig. 6e**; unlabeled pairs: 74.13 ± 17.84 %, IP-derived and OP-derived pairs: 130.50 ± 10.68 %; normalised to mean amplitude), indicating that individual PV interneurons connecting to both IP-derived and OP-derived neurons exhibit biases in connection strength to one of these neuron types over the other. These differences in the output of interneurons were not associated with differences in mean IPSC amplitude between the pair-types (**Supplementary Fig. 9**). Similarly, there were no differences in inter-somatic distances between pair types (**Fig. 6f**) or distance to pia (**Fig. 6g, Supplementary Fig. 9**). In conclusion therefore, individual interneurons can exhibit biased connectivity to L2/3 pyramidal neurons derived from different progenitor types, thereby defining the degree to which neighboring pyramidal neurons share local inhibitory input. This provides a lineage-based cellular mechanism for inhibitory subnetworks within L2/3.

**Figure 6:**
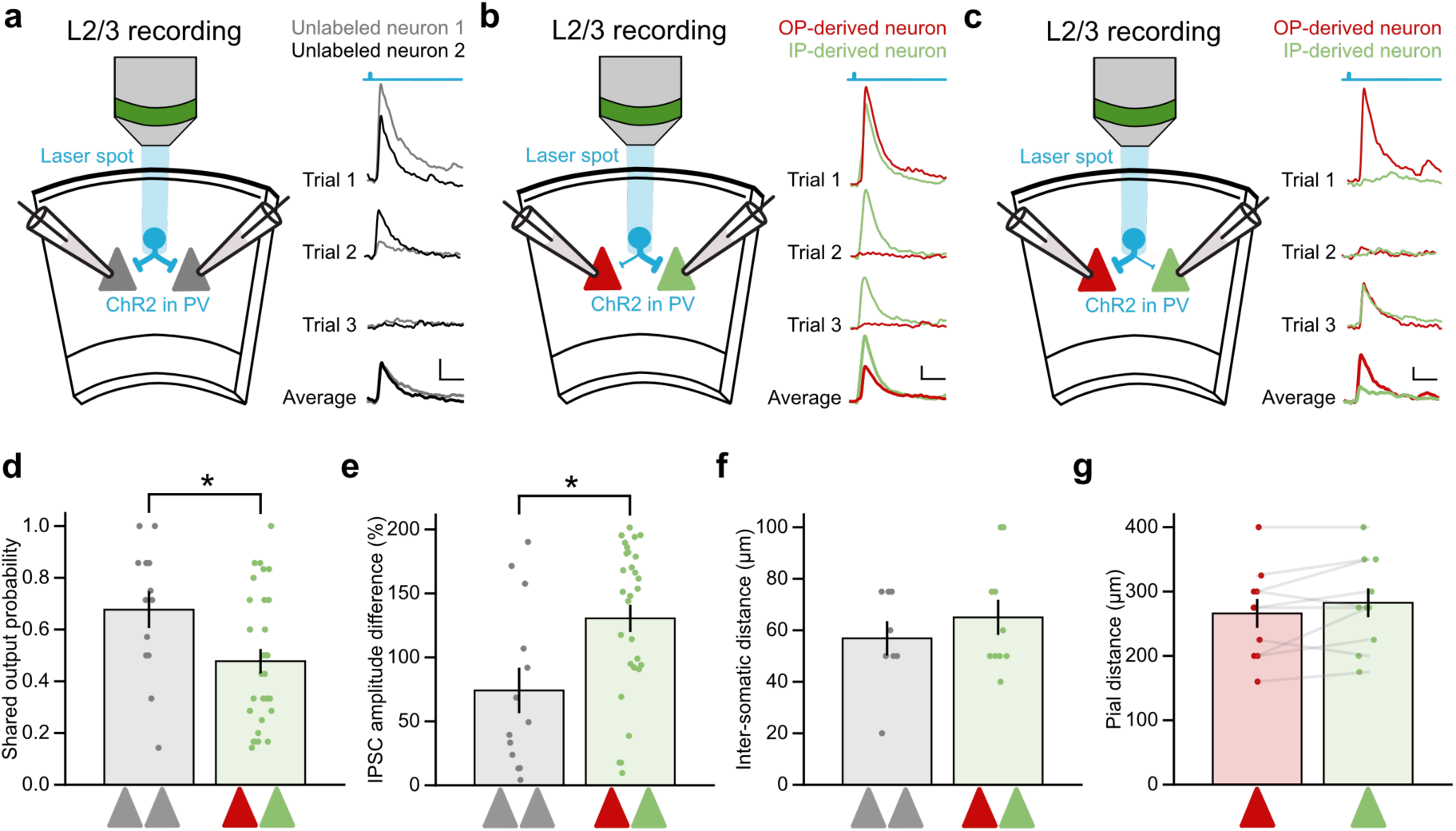
Interneurons differentially target L2/3 pyramidal neurons based on post-synaptic progenitor type. (**a**) Monosynaptic IPSCs were simultaneously recorded from pairs of unlabeled pyramidal neurons, whilst a laser was used to focally activate pre-synaptic ChR2-expressing PV interneurons (left). PV interneurons were minimally stimulated to produce a mix of successes and failures in post-synaptic responses. Example trials from a pair of unlabeled L2/3 pyramidal neurons (right). Scale bar, 50 pA, 25 ms. (**b**) As in ‘a’, but recordings were performed from IP-derived and OP-derived neuron pairs (left). Example trials where the IP-derived neuron received biased input (right). (**c**) As in ‘b’ (Ieft). Example trials where the OP-derived neuron received biased input (right). (**d**) The probability of observing a shared output was lower in IP-derived and OP-derived pairs, than for unlabeled pairs (two-tailed Mann-Whitney U test, *n* = 13 PV interneurons to 8 pairs of unlabeled neurons from 5 animals, *n* = 30 PV interneurons to 10 pairs of IP-derived and OP-derived pairs from 6 animals, *P* = 0.02, U = 278.5). (**e**) The IPSC amplitude difference for shared inhibitory inputs (expressed as percentage of mean amplitude) was greater for pairs comprising an IP-derived and OP-derived neuron, than for pairs of unlabeled neurons (two-tailed Mann-Whitney U test, *n* = 13 PV interneurons to 8 pairs of unlabeled neurons from 5 animals, *n* = 30 PV interneurons to 10 pairs of IP-derived and OP-derived pairs from 6 animals, *P* = 0.01, U = 102). (**f**) Inter-somatic distance was not different between pair types (two-tailed unpaired t-test, *n* = 8 pairs of unlabeled neurons from 5 animals, *n* = 10 pairs of IP-derived and OP-derived neurons from 6 animals, *P* = 0.41, T = - 0.84). (**g**) Distance to pia was not different between the IP-derived and OP-derived neurons (two-tailed paired t-test, *n* = 10 pairs from 6 animals, *P* = 0.61, df = 9, T = -0.52).

## Discussion

Synaptic inhibition underpins diverse computations within cortex, yet the principles by which inhibitory cortical circuits are organised remain poorly understood. Here we reveal that a pyramidal neuron’s synaptic inhibition depends upon the progenitor type from which the neuron is derived during embryonic development. By combining *in utero* fate mapping techniques with *in vivo* two-photon targeted patch-clamp recordings in the mature cortex, we show that progenitor type determines how a pyramidal neuron’s synaptic inhibition couples to local network activity. To determine the underlying synaptic mechanisms, we used optical circuit mapping to establish that these differences are not associated with the level of inhibition that pyramidal neurons receive from interneuron subclasses. Rather, progenitor type defines the degree to which pyramidal neurons share synaptic input from individual inhibitory interneurons, with interneurons differentially targeting pyramidal neurons based on post-synaptic progenitor type.

L2/3 pyramidal neurons exhibit sparse action potential firing *in vivo*, consistent with the fact that these neurons are under strong inhibitory synaptic control and that their post-synaptic GABAergic conductances exhibit tight temporal coupling with the LFP ^66–68^. Whilst earlier work characterised inhibitory cortical connectivity as non-specific ^6,7^, more recent evidence has shown that interneurons exhibit selectivity in the neurons that they target ^8–15^, and that individual pyramidal neurons can vary in the degree to which their inhibition is coupled to local network activity ^67^. Our *in vivo* patch clamp recordings reinforce the notion that neighboring excitatory neurons exhibit heterogeneity in their synaptic inhibition and we demonstrate that a source of this heterogeneity is the type of embryonic progenitor from which the pyramidal neuron is derived.

We are only beginning to understand how progenitor cell diversity shapes post-mitotic neuron diversity ^18,22,69–72^. Decades of research has established the concept that all pyramidal neurons are ultimately derived from multipotent RGCs, with neurons generated through direct asymmetric divisions or through indirect divisions via an IP ^27,73,74^. Therefore, the existence of multiple progenitor types enables neurons to be born simultaneously from distinct lineage trajectories. These different routes of excitatory neuron production may simply exist to amplify the number of neurons that can be produced during corticogenesis ^75,76^. Alternatively, each progenitor type may contribute to distinct wiring patterns in mature cortex ^34,37,77^. By revealing that different lineage trajectories contribute to diversity in local inhibitory circuitry, our results support the idea that multiple progenitor types diversify cortical circuits. This builds upon previous evidence that progenitor type influences excitatory synaptic connections in L2/3 ^34,78^ and strengthens the incentive to further explore progenitor subpopulations and their progeny.

The optical circuit mapping we performed suggests that regardless of their progenitor type, neighboring pyramidal neurons receive similar overall levels of synaptic inhibition from PV, SST and VIP interneurons. Hence, a L2/3 pyramidal neuron’s progenitor type does not appear to specify systematic differences in the input from major interneuron populations. As an alternative cellular mechanism, we considered the possibility that progenitor of origin determines the interactions between individual interneurons and individual pyramidal neurons. Firstly, using spontaneous currents as an unbiased way to sample inhibitory inputs, we revealed that progenitor type defines how much inhibitory input a pyramidal neuron shares with its neighbors. We then revealed the counterpart using optical minimal stimulation techniques, by showing that interneurons can differentially target neighboring pyramidal neurons based on the post-synaptic neuron’s progenitor type. These results constitute the first evidence that excitatory neuron progenitor type influences fine-scale inhibitory subnetworks in cortical circuits.

Our experiments do not resolve whether these subnetworks reflect the connectivity of interneuron subtypes that can be further subdivided beyond the single genetic markers we used here. These could include morphologically or molecularly defined interneuron subtypes ^79–81^, interneurons that reside in different cortical layers ^82–84^, or interneurons that are differentially recruited by long-range targets ^62^. Indeed, transcriptomic studies suggest that there are at least 10 different cortical PV interneuron subtypes ^79,80^, and it is possible that molecular mechanisms could specify the connections between a specific subtype of pre-synaptic interneuron and the post-synaptic pyramidal neuron, as has been proposed in L5 ^13^. It would therefore be interesting to explore in future work whether progenitor-defined populations of excitatory neurons preferentially receive certain types of inhibitory input across cortical layers or across transcriptomic inhibitory cell subtype. Meanwhile, recent work in hippocampus has shown that a neuron’s birthdate can influence the synaptic connectivity, morphology and synchrony of CA1 pyramidal neurons, including input and output connections with inhibitory interneurons ^85,86^. Further data from hippocampus has established that a clonal relationship, specifically excitatory sister neurons derived from the same individual progenitor, can affect the probability of shared inhibitory input ^87^. How progenitor type, birth dates and clonal relationships relate to one another remains to be determined, but the potential for these lineage trajectories to interact could generate yet further diversity within cortical circuits.

The lineage-related differences we have observed in inhibitory connectivity suggest that progenitor diversity may be responsible for establishing circuit motifs that differentially route information through cortex. The particular inhibitory inputs received by a pyramidal neuron has been shown to reflect the axonal target of that pyramidal neuron, consistent with the idea that local inhibitory subnetworks represent a mechanism for independently modulating different streams of excitatory information ^8–13,16^. In line with this framework, as well as engaging different local inhibitory circuits, IP-derived and OP-derived L2/3 pyramidal neurons have been shown to differ in their translaminar glutamatergic outputs to L5a and L5b, which are associated with cortical and subcortical information streams, respectively ^34,88,89^. It will therefore be interesting to establish whether progenitor diversity underlies a more general process for establishing excitatory-inhibitory cortical motifs, particularly given recent evidence that differences in long-range axonal patterns are related to whether a pyramidal neuron derives directly from a radial glial cell or an IP population ^90^, or indeed from different subpopulations of radial glial cells ^22,77^.

In summary, precise connectivity patterns between inhibitory and excitatory neurons underpin the coding properties of cortex, but how this circuitry is established is poorly understood. We demonstrate that L2/3 pyramidal neurons derived from intermediate progenitors receive inhibition that is less coupled to local network activity and that these neurons share less inhibitory inputs with neighboring neurons derived from other progenitors. This supports the idea that the construction of excitatory-inhibitory cortical subnetworks is embedded in embryonic development and that differences amongst progenitors contributes to diversity within these networks. We suggest that progenitor diversity may have evolved in part to generate heterogeneity in cortical circuits, and that progenitor identity may be an effective approach for understanding the origins of inhibitory subnetworks in different systems.

## Author contributions

G.G. and C.J.A. designed the study. G.G. conducted the experiments, wrote software and performed analysis. S.E.N. generated molecular tools. S.T. assisted in the collection and analysis of the *in vivo* LFP data. M.J.B., K.M. and R.J.B. provided mentorship and teaching of the techniques. G.G. and C.J.A. wrote the paper, with input from the other authors.

## Acknowledgements

We would like to thank members of the Akerman lab for advice and comments. The research leading to these results has received funding from the European Research Council under grant agreement number 617670; plus BBSRC project BB/S007938/1. In addition, G.G. was supported by a Wellcome Trust Doctoral Fellowship, S.E.N. was supported by a Royal Society Dorothy Hodgkin Fellowship, S.T. was supported by the Rhodes Trust, M.J.B. was supported by a University of Oxford Clarendon Scholarship, R.J.B. was supported by a Shaun Johnson Memorial Scholarship sponsored by the Leverhulme Trust and Mandela Rhodes Foundation.

## Code Availability

Analysis scripts will be made available upon request.

## Methods

### Experimental animals

Experiments were performed using both male and female mice, which were bred, housed, and used in accordance with the UK animals (Scientific Procedures) Act (1986). Animals were reared in a dedicated facility with a 12-hour light/dark cycle with access to food and water ad libitum. Unless stated, experiments were conducted in C57BL/6 wildtype mice. For optogenetic experiments, mice were the result of breeding crosses between a homozygous *floxed* ChR2 mouse (B6;129S-*Gt(ROSA)26Sor^tm^*^32*(CAG-COP4*H134R/EYFP)Hze*^/J) and either a homozygous PV-Cre mouse (B6;129P2-Pvalb^tm1(cre)Arbr^/J), a homozygous SST-Cre mouse (Sst^tm2.1(cre)Zjh^/J), or a homozygous VIP-Cre mouse (Vip^tm1(cre)Zjh^/J). Mice were purchased from The Jackson Laboratory (USA).

### *In utero* electroporation

Females were paired for timed matings and checked daily for vaginal plugs. Embryonic day 0.5 (E0.5) was defined as midday on the day that the vaginal plug was detected. *In utero* electroporation was performed at E15.5 using standard procedures. Dams were induced and maintained under anaesthesia with isofluorane (Zoetis). Depth of anaesthesia was assessed by monitoring the animal’s respiration rate and righting reflex. Surgery was performed under sterile conditions and dams were operated on a heating mat to prevent loss of body temperature. Viscotears (Bausch and Lomb) was applied to protect the corneas during surgery. Meloxicam (5 mg/kg) (Boehringer Ingelheim Vetmedica Gmbh) and Buprenorphine (0.1 mg/kg) (Vetergesic Multidose, Alstoe Animal Health) analgesics were administered by subcutaneous injection. The abdomen was shaved and sterilized using Hibiscrub (Regent Medical). A midline laparotomy was performed and the uterine horns were exposed. A mixture of plasmid DNA (∼1 µg/µl) and 0.03 % fast green dye (Sigma-Aldrich) was injected intraventricularly in each embryo through the uterine wall and amniotic sac, with a glass micropipette (World Precision Instruments; outer diameter (OD): 1.5 mm, inner diameter (ID): 1.12 mm) that had been pulled using a Flaming/Brown Micropipette Puller (Sutter Instrument Company). Plasmids were prepared using an EndoFree plasmid kit (Qiagen). The total volume injected per embryo was ∼1.5 µL. All plasmids were injected as a 1:1 ratio.

Plasmid DNA included:

i. ‘pTα1-Cre’, in which Cre recombinase is under the control of a portion of the Tubulin alpha-1 (Tα1) promoter (Tα1-Cre was a gift from Tarik Haydar) ^51^.
ii. ‘pTα1-Flpo’, in which codon optimized Flp recombinase is under the control of a portion of the Tubulin alpha-1 (Tα1) promoter. To generate Tα1-Flpo, Cre recombinase and its associated 3’ poly(A) sequences were removed from Tα1-Cre using XhoI/SacI and replaced with Flpo recombinase and its associated poly(A) sequences from pCAG-Flpo (a gift from Massimo Scanziani Addgene plasmid # 60662; http://net.addgene:60662; RRID;Addgene 60662). The Flpo sequence was amplified using Phusion DNA polymerase (New England Biolabs) and the primers Flpo_XhoI_Fwr gagaagctcgaggccgccaccatggctcctaaga and Flpe_pATerm_SacI_Rev (ctgaatgagctcgggctgcaggtcgagggatct). The PCR product was digested and ligated into the Tα1 backbone.
iii. ‘pCAG-FLEX-GFP’, a single colour Cre-dependent reporter that uses the chicken β-actin (CAG) promoter to control a flexible excision (FLEx) cassette, whereby Cre recombination permanently turns on expression of enhanced green fluorescent protein (GFP). pCAG-FLEX-GFP was a gift from Edward Boyden (Addgene plasmid # 28304; http://n2t.net/addgene:28304; RRID:Addgene_28304).
iv. ‘pEf1a-FRT-tdTom/GFP’, a dual colour Flp-dependent reporter that uses the human elongation factor-1a promoter (Ef1a) to express tdTomato, until Flp recombination permanently switches expression to GFP. To generate pEf1a-FRT-tdTom/EGFP, ChR2-EYFP was removed from the backbone vector pAAV-Ef1a-fDIO-hChR2(H134R)-EYFP using AscI/NheI and replaced with tdTomato/EGFP from pAAV-Ef1a-DO_DIO-TdTomato_EGFP-WPRE-pA digested using the same restriction enzymes. pAAV-Ef1a-fDIO-hChR2(H134R)-EYFP was a gift from Karl Deisseroth (Addgene plasmid # 55639; http://n2t.net/addgene:55639; RRID:Addgene_55639). pAAV-Ef1a-DO_DIO-TdTomato_EGFP-WPRE-pA was a gift from Bernardo Sabatini (Addgene plasmid # 37120; http://n2t.net/addgene:37120; RRID:Addgene_37120).
v. “pTbr2-Flpo”, in which Flp recombinase is under the control of a portion of the T-box brain protein 2 (Tbr2) promoter. To generate pTbr2-Flpo, first pTbr2-GFP was generated in the retroviral backbone pCAG-GFP (CAG-GFP was a gift from Fred Gage (Addgene plasmid # 16664 ; http://n2t.net/addgene:16664 ; RRID:Addgene_16664). The CAG promoter was removed using restriction enzymes SalI/AgeI and replaced with a 2.5kb fragment of the mouse Tbr2 promoter based on previous work ^91^. The promoter region was amplified from mouse brain genomic DNA using the following primers:

MuTbr2FwrS_Sal1: CTGCAGAAGTCGACTTTACTGAGGTGGGGTTCCAG

MuTbr2Rev_AgeI: TTCTGCAGACCGGTGCTTTAGCGAATCGCAGACG

The GFP sequence was subsequently removed and replaced with a small multiple cloning site (MCS) containing sites AgeI, HindIII, StuI and BamHI inserted between the Tbr2 promoter and the improved Cre recombinase sequence (iCre) to generate pTbr2-Cre. iCre was amplified from pAAV-CAG-iCre (pAAV-CAG-iCre was a gift from Jinhyun Kim (Addgene plasmid # 51904 ; http://n2t.net/addgene:51904 ; RRID:Addgene_51904). Subsequently, pTbr2-Flpo was generated by removing the iCre sequence using a BamHI/PmeI digest and replacing it with the Flpo sequence amplified from pCAG-Flpo (pCAG-Flpo was a gift from Massimo Scanziani (Addgene plasmid # 60662 ; http://n2t.net/addgene:60662 ; RRID:Addgene_60662) using the following primers:

BamHI-Flpo-Fwr:

CTGCAGAAGGATTCGCCGCCACCATGGCTCCTAAGAAGAAGAGGA PmeI-Flpo-Rev: ATGACGTCGTTTAAACTCAGATCCGCCTGTTGATGTAG

(vi) “pGlast-Cre”, in which Cre recombinase is under the control of a portion of the glial high affinity glutamate transporter (GLAST) promoter and was a gift from Tarik Haydar^44^.
(vii) “pCAG-FRT-TdTom”, a single-colour Flpo-dependent reporter that uses the CAG promoter to control a flp-dependent double-floxed inverted open-reading frame, whereby Flpo expression permanently switches on the expression of tdtomato. pCAG-FRT-TdTom was derived from pAAV-CAG-fDIO-mNeonGreen (pAAV-CAG-fDIO-mNeonGreen was a gift from Viviana Gradinaru (Addgene plasmid # 99133 ; http://n2t.net/addgene:99133 ; RRID:Addgene_99133). The mNeonGreen sequence was removed using a AscI/NheI digest and replaced with a Tandem Tomato (TdTomato) sequence that was amplified from pCAG-mNaChBac-T2A-tdTomato (pCAG-mNaChBac-T2A-tdTomato was a gift from Massimo Scanziani (Addgene plasmid # 60650 ; http://n2t.net/addgene:60650 ; RRID:Addgene_60650) using the following primers to amplify both fluorescent protein sequences:
AscI-eGFPRev-Fwr: GAGAACGGCGCGCCTTACTTGTACAGCTCGTCCATGC
NheI-eGFPFwr-Rev: GAGACCGCTAGCGCCACCATGGTGAGCAAGG
(viii) “pCAG-FRT-GFP”, a single-colour Flpo-dependent reporter that uses the CAG promoter to control a flp-dependent double-floxed inverted open-reading frame, whereby Flpo expression permanently switches on the expression of GFP. CAG-FRT-GFP was derived from pAAV-CAG-fDIO-mNeonGreen by replacing the mNeonGreen with GFP from pCAG-GFP.

Electroporation was performed by placing the anode of the platinum Tweezertrodes (Genetronics) over the dorsal telencephalon. Five pulses (50 ms duration separated by 950 ms) at 40 V were delivered with a BTX ECM 830 square pulse generator (Genetronics). All accessible embryos underwent electroporation. The uterus was lavaged with warmed, sterile phosphate-buffered saline (PBS) and replaced into the abdomen. The abdominal muscle and skin incision were closed with Vicryl and Prolene (Ethicon) sutures, respectively. Animals were recovered from anaesthesia in a chamber heated to 38 °C. Meloxicam and Buprenorphine were provided orally during the postoperative window, as appropriate. Dams were allowed to litter down naturally, and the day of birth was recorded as postnatal day 0 (P0).

### *In vivo* electrophysiology

Animals (aged 4-6 weeks old) were anaesthetized and maintained with 0.8-1.5% isoflurane for the duration of the recording. Depth of anaesthesia was assessed using breathing rate and toe pinch reflex. Animals were secured into the ear bars (Narishige), placed on a heat pad to maintain body temperature and Viscotears was applied to protect the corneas. The hair over the scalp was shaved and treated with hair removal cream (Veet). Craniotomies were performed under a stereoscope (Olympus) and were targeted over barrel cortex (coordinates: -1 mm AP, +3 mm ML, from bregma). An incision was made to expose the cranium and the surrounding skin was fixed with superglue (Loctite). Dental cement (Simplex Rapid) was used to build a chamber around the exposed skull, which could hold 1-2 drops of cortex buffer. Successive rounds of drilling were made using a dental drill (Foredom) to gently loosen the cranium, whilst the cranium was kept moist with cortex buffer, containing (in mM): 125 NaCl, 5 KCl, 10 HEPES, 2 MgSO_4_·7H_2_O, 2 CaCl_2_·2H_2_O, 10 Glucose. The skull and underlying dura were carefully removed with a 0.3 mL syringe needle. The brain was submerged with cortex buffer throughout the recording. A silver-chloride ground electrode (Multichannel systems) was placed on the skull anterior to the craniotomy.

The animal was transferred to an upright microscope (Olympus BX51WI) that had been customized for *in vivo* recordings. Pyramidal neurons in L2/3 of S1 were targeted using methods described previously ^92–94^. Briefly, a cesium-based internal solution (containing, in mM: 126 CsOH.H_2_0, 126 gluconic acid, 10 HEPES, 5 TEA-Cl, 2 QX-314 Cl, 4 Mg_2_ATP, 0.4, NaGTP, 10 Na_2_ Phosphocreatine, 2 EGTA, 0.2 % Biocytin, 0.02 Alexa594, pH 7.2-7.3, 290-295 mOsmol/L) was used to fill micropipettes with a tip resistance of 5-8 MΩ, pulled from borosilicate glass (Harvard Aparatus; OD: 1.5 mm, ID: 0.86 mm) using a vertical puller (Narishige). Positive pressure (800 mbar) was applied whilst the pipette was placed onto the surface of the brain and located using a 4x objective (Olympus PLN; NA: 0.1). Upon entering cortex by 80 µm, the positive pressure was reduced to 70 mbar and the pipette was advanced in 10 µm/s increments (up to a maximum depth of 350 µm) whilst looking for large changes in resistance, as an indicator of the close proximity of a cell membrane. At this point, the positive pressure was released to facilitate gigaseal formation and after approximately 1 minute in gigaseal, the whole-cell configuration was achieved by applying small suction pulses to the pipette. GFP-expressing neurons were targeted using a custom-built two-photon microscope, equipped with a 40 x water-immersion objective (Olympus LUMPLFLN, NA: 0.8), modified confocal scan unit (Olympus, FV300) and Ti:Sapphire laser (Newport Spectra-Physics Mai Tai HP; wavelength: 850 nm, power: 50 mW). The mean cell distance from pia was 111.59 ± 8.60 µm and did not differ between experimental conditions (two-tailed unpaired t-test, unlabeled neuron mean: 123.27 ± 14.11 µm, *n* = 11 neurons from 5 animals, IP-derived neuron mean: 99.91 ± 9.18 µm, *n* = 11 neurons from 5 animals, *P* = 0.18, df = 20, T = 1.39).

Inhibitory post-synaptic currents (IPSCs) were recorded in voltage clamp mode at a holding potential of +20 mV, close to the reversal potential of glutamate receptors. This led to the detection of outward synaptic currents, consistent with a strong driving force on the GABA_A_ receptor ^52^. LFP recordings were made using a glass pipette that had been broken to a resistance of 0.5-1 MΩ. LFP pipettes were slowly inserted into the brain, to be in close proximity to the patch pipette. Electrophysiological data were acquired using a Multiclamp 700B amplifier (Molecular Devices) and digitizer (Digidata 1550, Molecular Devices), controlled by Clampex software (Molecular Devices). The LFP signal was band passed filtered at 0.1 – 200 Hz and then digitized at 10 kHz. Spontaneous IPSCs and LFP activity were recorded for 7.5 minutes. Every 15 s, a 10 mV hyperpolarizing pulse (100 ms duration) was applied to the command voltage of the patch pipette to monitor access resistance. Mean access resistance across all cells was 58.48 ± 4.23 MΩ. To account for the effects of access resistance, offline correction was performed ^95^. This involved multiplying the measured current response with 90 % of the access resistance, and then subtracting this from the command voltage to produce a corrected holding potential. Current traces were converted into inhibitory synaptic conductances by using the driving force calculated from the difference between the estimated chloride equilibrium potential (-73 mV) and the corrected holding potential.

Inhibitory synaptic inputs were detected using the first derivative of the conductance trace from the patched neuron. Firstly, conductance traces were median-subtracted, low-pass filtered at 100 Hz, and down-sampled to 4 kHz. Peaks in the first derivative were identified using a threshold-based method. Input start and input peak were defined as times at which the peak in the first derivate returned to 10% of its peak value before and after its maximum, respectively. Input amplitudes were then calculated by subtracting the conductance value at the input start time from the conductance value at the input peak time. Input amplitude threshold was set at 40 pA to maximise the detection of action-potential-evoked inhibitory conductances ^59,64^.

LFP traces were median-subtracted and down-sampled to 4 kHz to match the conductance traces. The input-triggered LFP (IT-LFP) was calculated by averaging all 100 ms LFP epochs that were centred on the start time of the defined inhibitory synaptic inputs detected in the patched neuron. The peak of the IT-LFP was defined as the mean IT-LFP during the 10 ms following the onset of the inhibitory input. LFP events were identified by detecting peaks in the LFP recording that were at least 100 µV in amplitude and with a duration of 50 ms at 50 % peak height. An event time tiling coefficient (ETTC) was calculated from the LFP events and the inhibitory synaptic inputs/events using methods described previously ^96–98^. Briefly, this measure calculates the proportion of coincident events between two traces (traces A and B) that occur above chance level, by calculating the proportion of events in trace A that fall within a short temporal window (± dt) of events in trace B. We used a dt of 25 ms for our analysis. To account for events that are expected to co-occur by chance, the proportion of trace A that occurs within ± dt windows from events in trace A is subtracted. The measure is then calculated reciprocally for trace B with respect to events in trace A. This generates a coefficient that reflects trace A’s relationship to trace B, and a coefficient that reflects trace B’s relationship to trace A. The mean of the two coefficients is the ETTC, and therefore represents a bi-directional measure of synchrony.

### *In vitro* electrophysiology

Acute cortical slices were prepared from primary somatosensory cortex (S1). Animals (aged 3-6 weeks old) were anaesthetized with isoflurane and then decapitated. Coronal 350 µm slices were cut using a vibrating microtome (Microm HM650V). Slices were prepared in artificial cerebrospinal fluid (aCSF) containing (in mM): 65 Sucrose, 85 NaCl, 2.5 KCl, 1.25 NaH_2_PO_4_, 7 MgCl_2_, 0.5 CaCl_2_, 25 NaHCO_3_ and 10 glucose, pH 7.2-7.4, bubbled with carbogen gas (95% O_2_ / 5% CO_2_). Slices were immediately transferred to a storage chamber containing aCSF (in mM): 130 NaCl, 3.5 KCl, 1.2 NaH_2_PO_4_, 2 MgCl_2_, 2 CaCl_2_, 24 NaHCO_3_ and 10 glucose, pH 7.2-7.4 and bubbled with carbogen gas for 1 hour at 35 °C. When required, slices were transferred to the glass coverslip that formed the bottom of the recording chamber, and continuously perfused with aCSF bubbled with carbogen gas (perfusion speed 3 ml/min).

Whole-cell patch-clamp recordings were performed using glass pipettes (4-6 MΩ), pulled from standard wall borosilicate glass capillaries (Harvard Apparatus; OD: 1.2 mm ID: 0.69 mm) and containing (in mM): 126 CsOH.H_2_0, 126 Gluconic acid, 10 HEPES, 1.5 MgCl_2_.6H_2_0, 2 QX-314Cl, 4 Mg_2_ATP, 0.4 NaGTP, 10 Na Phosphocreatine, 2 EGTA, (pH 7.2-7.3, osmolarity, 290-295 mOsmol/l) and 0.25 mg/ml biocytin. Pairs of neighboring L2/3 pyramidal neurons approximately 40 µm below the surface of the slice were visualized for simultaneous patch-clamp recording under an upright microscope (Olympus BX51WI) equipped with a 20x water-immersion objective (Olympus XLUMPLFLN; 1.0 NA), dodt gradient contrast, and epifluorescence illumination. IP-derived and OP-derived L2/3 pyramidal neurons were identified using epifluorescence and appropriate filter sets for separating GFP and TdTomato signals.

*In vitro* electrophysiological data were acquired using a Multiclamp 700A amplifier (Molecular Devices) and digitizer (ITC-18, Instrutech), controlled by WinWCP software (University of Strathclyde, UK, RRID:SCR014713). Access resistance was measured during every sweep on all experimental protocols using a hyperpolarizing pulse (10 mV, 100 ms duration), and only recordings where access resistance was less than 30 MΩ were included. Access resistance did not systemically vary between any of the experimental groups. To detect spontaneous action-potential-evoked IPSCs (sIPSCs), 7.5 minutes of spontaneous activity was recorded whilst holding each neuron at 0 mV, at the reversal potential of glutamate receptors under these recording conditions. sIPSCs were observed as large outward currents and were identified using the same detection method described for *in vivo* inhibitory conductances (see above). Traces were low-pass filtered at 100 Hz and down-sampled to 4 kHz. To maximize the detection of action-potential-evoked IPSCs, an amplitude threshold of 40 pA was applied, which was selected based on reported values of unitary IPSCs ^59,64,99^. However, the same effects were seen across a range of amplitude thresholds (**Supplementary Figure 5C**).

For widefield optogenetic stimulation of ChR2-expressing interneurons, blue light pulses (473 nm, 1 ms duration, 0.05 Hz) were delivered via a high-power light emitting diode (LED; Luxeon Star, Farnell) positioned below the recording chamber, so as to illuminate the entire brain slice. Light intensity recorded at the tissue surface closest to the LED was 10-40 mW/mm^2^. A different optical setup was used to activate putative single ChR2-expressing PV interneurons, which involved delivering a spot of blue light to the upper surface of the brain slice (40 µm diameter at the slice surface, 1 ms duration, 0.05 Hz) from a diode-pumped solid-state laser (473 nm peak wavelength; Shanghai Laser and Optics Century). The laser was coupled to a 200 μm diameter multimode optic fiber via a collimating lens (Thorlabs), the tip of the optic fiber was positioned at an image plane in the center of the microscope’s optical axis, and the light was directed into the objective (Olympus XLUMPLFLN; 20 x, 1.0 NA) via a dichroic mirror. In line with previous work that has used minimal optogenetic stimulation to reveal shared connections ^100^, we adjusted the intensity of the light spot (0.3–4.2 mW/mm^2^) to a regime where we observed post-synaptic IPSC successes and failures, analogous to electrical minimal stimulation conditions. When separate PV interneurons were stimulated for the same pair of pyramidal neurons, the stimulation sites were on average 611.01 ± 90.62 µm apart. If the axon of the stimulated PV interneuron connected with similar probability and strength to the two post-synaptic pyramidal neurons, the optically-evoked IPSCs would be expected to co-fluctuate across stimulations, exhibiting similar probability and amplitude on a trial-to-trial basis.

### Post-hoc histology

Following *in vivo* electrophysiology, animals were fixed by cardiac perfusion with 4% PFA (Sigma Aldrich) in 0.1 M PBS (pH 7.4) and then left in PFA for an additional 24-48 hours. Fixed brains were then sectioned at 100 µm on a vibratome (Microm). Following *in vitro* electrophysiology, slices were fixed in 4% PFA in 0.1 M PBS (pH 7.4) for 24 hours then transferred to 0.1 M PBS. To visualize biocytin-filled neurons, sections were incubated in streptavidin Alexa Fluor 647 (1:1000, Thermo Fisher) for 2 hours at room temperature. Sections were then stained with 4’,6-Diamidino-2-Phenylindole, Dihydrochloride (DAPI, 1:10,000, Thermo Fisher). Three PBS washes were performed in between each step. Sections were then mounted with Vectashield (Vectorlabs) onto glass slides (Avantor) and sealed. Images were acquired with an LSM 710 confocal microscope using ZEN software (Zeiss).

### Data analysis and statistical tests

Data were assessed for normality and appropriate parametric (1 sample t-test, paired t-test or unpaired t-test) or non-parametric (Wilcoxon signed rank test, Mann-Whitney U test) statistical tests were applied. For tests that had more than 2 groups, either a one-way ANOVA (parametric data) or Kruskal Wallis (non-parametric data) was used to test for differences between groups. If differences were detected, Bonferroni’s correction was used to adjust *P* values for multiple comparisons. Statistically significant differences are indicated with asterisks on graphs as * *P* < 0.05, ** *P* < 0.01, *** *P* < 0.001. Analysis was carried out using custom written scripts in Python 3.7. Data are presented as mean ± sem unless otherwise stated.

## Supplementary Figures

**Supplementary Figure 1:**
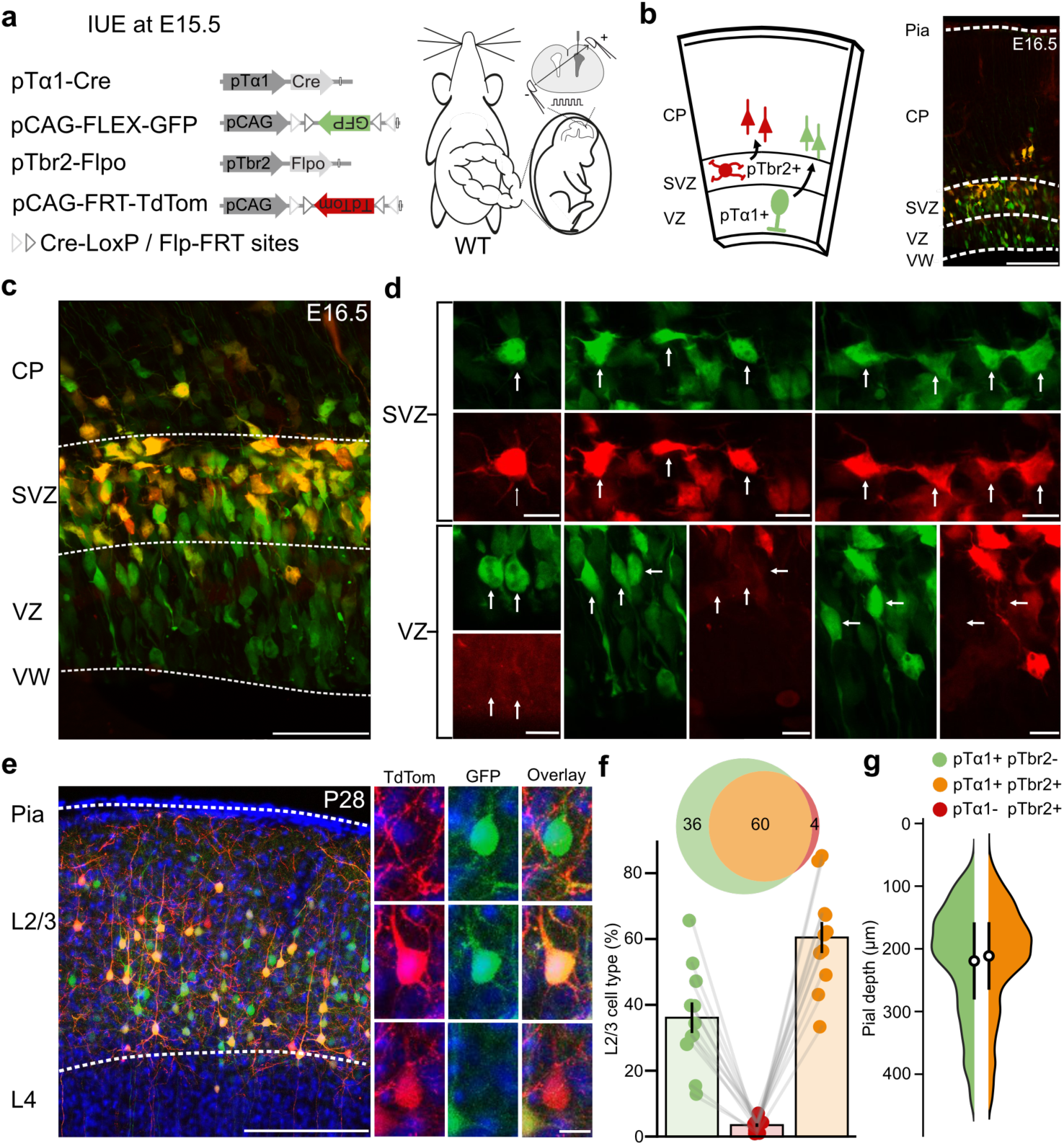
Intermediate progenitors labelled by Tα1 promoter activity comprise a population of apical IPs and basal IPs. (**a**) Four plasmid IUE of Tα1-Cre, Cre-dependent GFP, Tbr2-Flpo and Flpo-dependent TdTomato was performed at E15.5 to label intermediate progenitors and their progeny. (**b**) Tissue was collected 24 hours after IUE at E16.5 to visualise IPs and newly born IP-derived neurons in the VZ and SVZ. Scale bar, 100 µm. (**c**) Tα1-positive/Tbr2-negative (i.e. expressing only GFP) aIP-derived neurons can be seen in the ventricular zone (VZ) and Tα1-positive/Tbr2-positive (i.e. expressing GFP and TdTom) bIP-derived neurons can be seen in the subventricular zone (SVZ). Scale bar 50 µm. VW = ventricular wall, CP = cortical plate. (**d**) Selected enlarged sections from **‘c’**. GFP and TdTomato channels for each image are shown separately. The top row shows examples of multipolar Tα1-positive/Tbr2-positive bIPs residing in the SVZ. The bottom row shows examples of Tα1-positive/Tbr2-negative aIPs with radial morphology in the VZ, some of which are dividing. Scale bars, 20 µm. (**e**) L2/3 IP-derived neurons at P28 following IUE of the plasmids in ‘**a**’. Scale bar, 100 µm. Insets (right) show examples of neurons derived from a Tα1-positive/Tbr2-negative lineage (top), a Tα1-positive/Tbr2-positive lineage (middle) and a Tα1-negative/Tbr2-positive lineage (bottom). Scale bar, 20 µm. (**f**) Percentage of each L2/3 cell type. Tα1-positive/Tbr2-negative mean: 36.08 ± 4.66 %, Tα1-positive/Tbr2-negative mean: 60.46 ± 4.73 %, Tα1-negative/Tbr2-positive mean: 3.46 ± 0.55 %, n = 11 brains. (**g**) Tα1-positive/Tbr2-negative and Tα1-positive/Tbr2-positive lineage sat at equal distances from pia (two-tailed Wilcoxon signed-rank test, Tα1-positive/Tbr2-negative mean: 219.34 ± 18.73 µm, Tα1-positive/Tbr2-positive mean: 211.37 ± 16.31 µm, *n* = 11 brains, *P* = 0.24, W = 19). Pial depth is shown as a kernel density estimate alongside mean ± standard deviation.

**Supplementary Figure 2:**
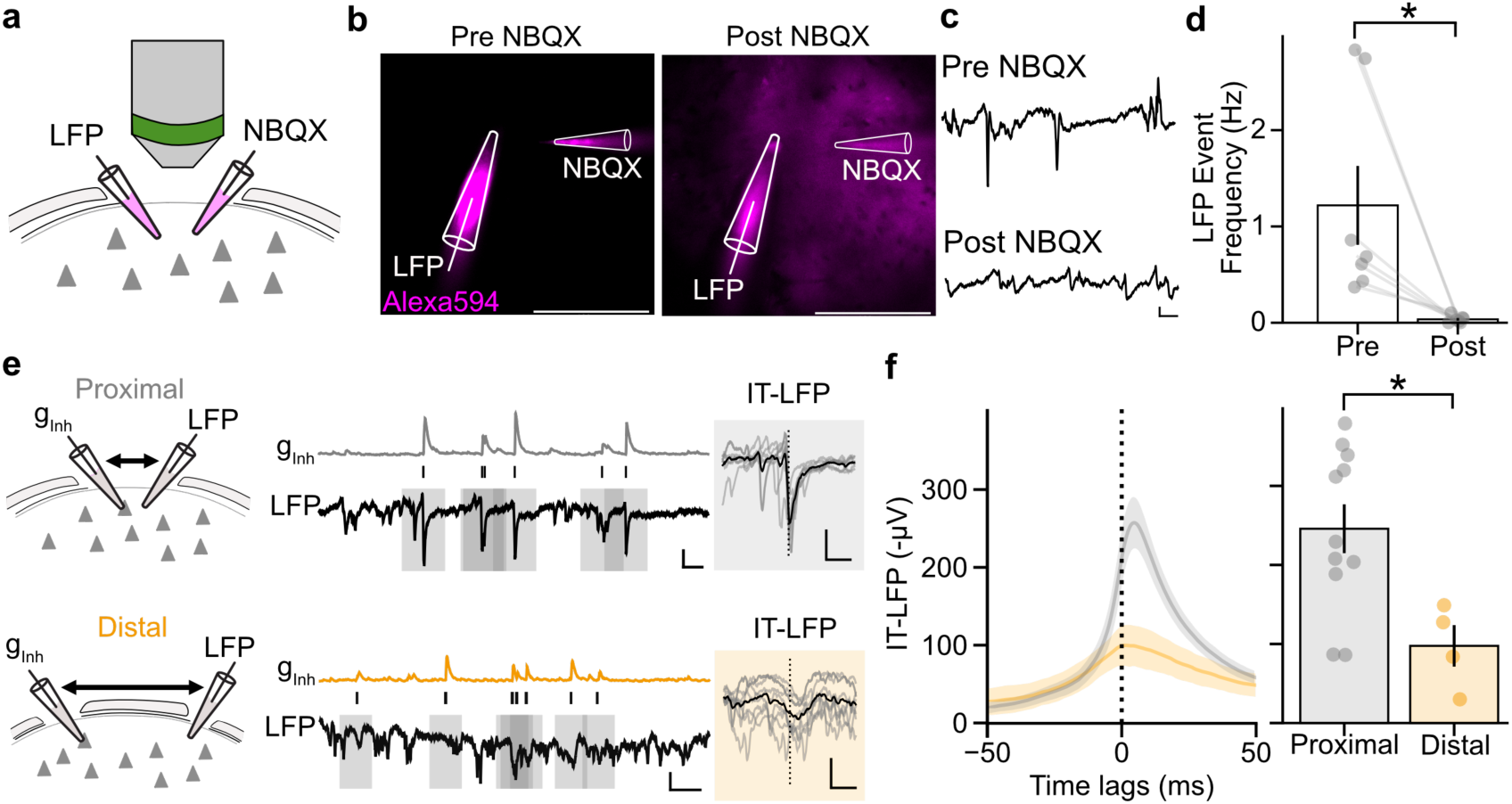
LFP reflects local synaptic activity and can be used as a coupling measure. (**a**) LFP was recorded before and after a local injection of the AMPA receptor antagonist NBQX *in vivo*. (**b**) Alexa594 was added to the NBQX pipette such that successful NBQX injection was confirmed through visualisation of Alexa594 in the extracellular space. Scale bar, 50 µm. (**c**) Example traces showing sharp deflections in LFP activity, reflecting nearby synaptic potentials that were blocked by NBQX. Scale bar 50 µv, 250ms. (**d**) LFP event frequency was significantly reduced following NBQX injection (two-tailed Wilcoxon test, pre-NBQX mean: 1.22 ± 0.41 Hz, post-NBQX mean: 0.04 ± 0.01 Hz, n = 7 recordings from 7 animals, P = 0.016, W = 0). (**e**) Simultaneous LFP and whole-cell patch clamp recordings were performed proximally to one another, under 500 µm apart. LFP and ginh were tightly correlated on a millisecond timescale, reflected in the large input-triggered LFP (IT-LFP) (top). Scale bar 2 nS, 200 µV, 250 ms, IT-LFP scale bar 200 µV, 100 ms. Recordings were also performed distally, where the LFP and patch pipette were in separate craniotomies separated by more than 1 mm. LFP and ginh were less tightly coupled, reflected in the smaller IT-LFP (bottom). Scale bar 2 nS, 200 µV, 250 ms, IT-LFP scale bar 200 µV, 100 ms. (**f**) IT-LFP was significantly larger in proximal recordings than distal recordings (two-tailed unpaired t-test, proximal mean: 246.27 ± 31.00 µV, n = 11 neurons from 5 animals, distal mean: 97.79 ± 26.30 µV, n = 4 neurons from 3 animals, *P* = 0.017, T = 2.73).

**Supplementary Figure 3:**
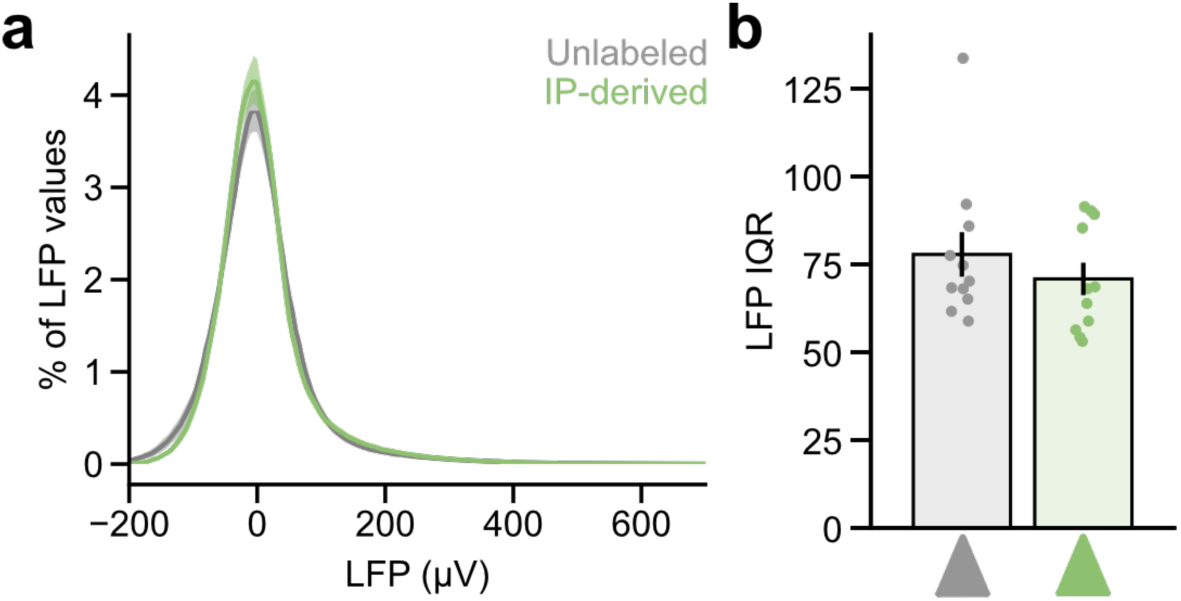
The distribution of LFP values is similar across unlabeled and IP-derived *in vivo* recordings. (**a**) Distribution of LFP values for unlabeled and IP-derived recordings. (**b**) The interquartile range (IQR) for LFP values did not differ between unlabeled and IP-derived recordings (two-tailed Mann-Whitney U test, unlabeled mean: 78.11 ± 6.2, *n* = 11 recordings from 5 animals, IP-derived mean: 70.77 ± 4.63, *n* = 11 recordings from 5 animals, *P* = 0.36, U = 75).

**Supplementary Figure 4:**
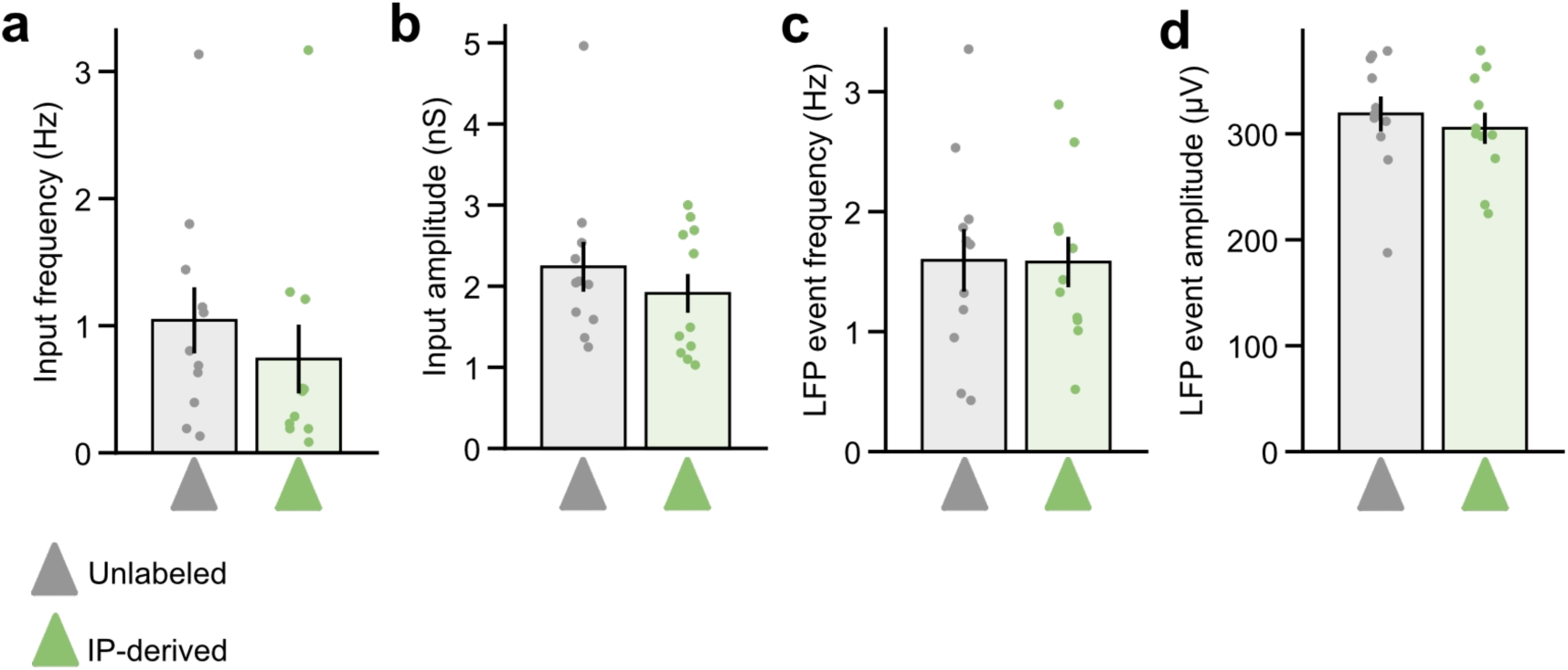
Inhibitory synaptic inputs and LFP events are similar across unlabeled and IP-derived *in vivo* recordings. (**a**) Inhibitory input frequency did not differ between unlabeled and IP-derived neurons (two-tailed Mann-Whitney U test, unlabeled mean: 1.13 ± 0.25 Hz, *n* = 11 neurons from 5 animals, IP-derived mean: 0.83 ± 0.29 Hz, *n* = 11 neurons from 5 animals, *P* = 0.21, U = 80). (**b**) Inhibitory input amplitude did not differ between unlabeled and IP-derived neurons (two-tailed Mann-Whitney U test, unlabeled mean: 2.24 ± 0.31 nS, IP-derived mean: 1.91 ± 0.24 nS, *P* = 0.55, U = 70). (**c**) LFP event frequency did not differ between recordings from unlabeled and IP-derived neurons (two-tailed unpaired t-test, unlabeled mean: 1.57 ± 0.24 Hz, *n* = 11 recordings from 5 animals, IP-derived mean: 1.58 ± 0.22 Hz, *n* = 11 recordings from 5 animals, *P* = 0.96, df = 20, T = -0.45). (**d**) LFP event amplitude did not differ between recordings from unlabeled and IP-derived neurons (two-tailed Mann-Whitney U test, unlabeled mean: 319.30 ± 16.41 µV, *n* = 11 recordings from 5 animals, IP-derived mean: 305.17 ± 14.80 µV, *n* = 11 recordings from 5 animals, *P* = 0.53, df = 20, U = 0.64).

**Supplementary Figure 5:**
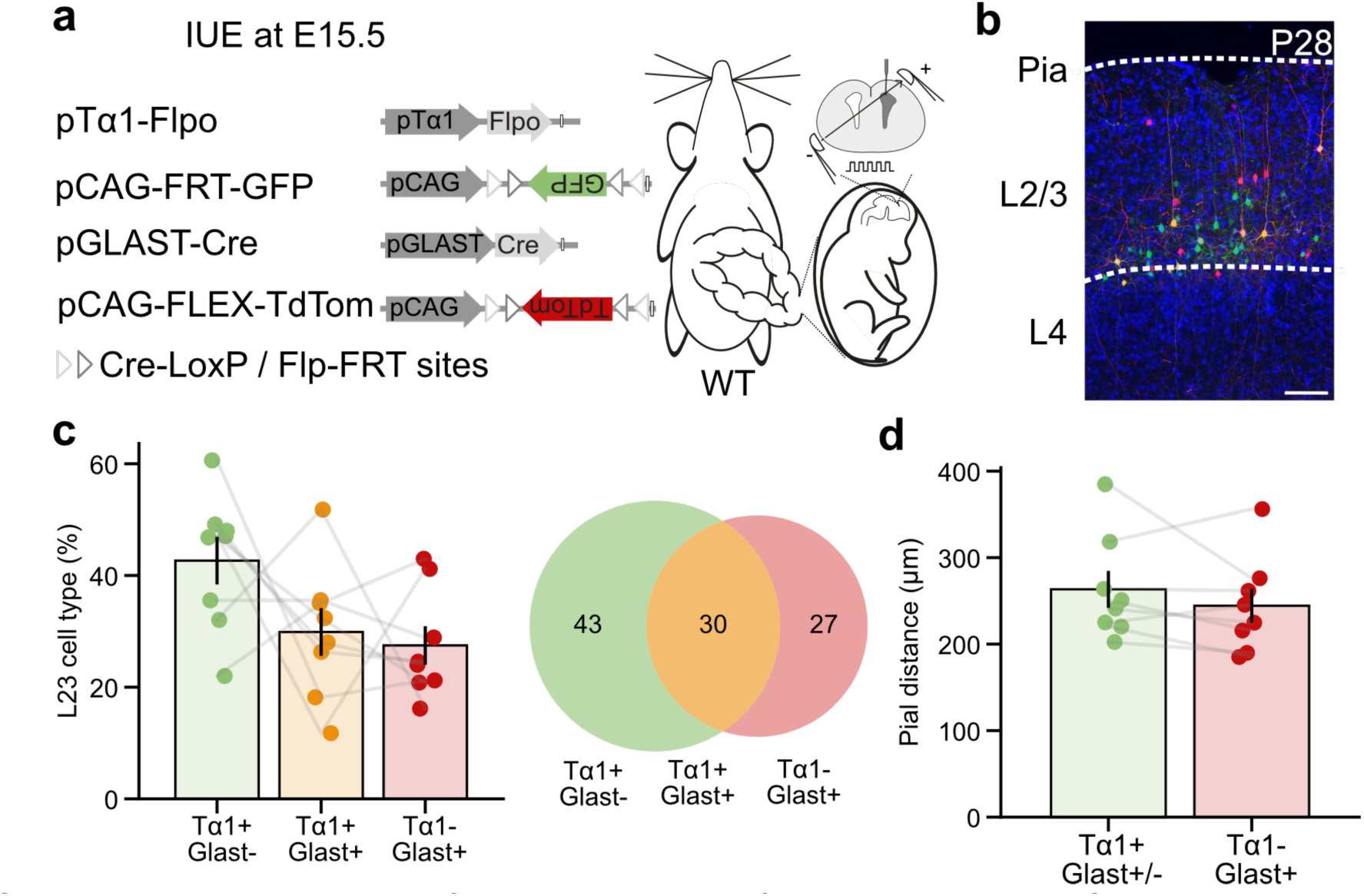
Characterization of neurons derived from a Tα1-negative lineage. (**a**) Animals underwent a four plasmid IUE of pTα1-Flpo, pCAG-FRT-GFP, pGlast-Cre and pCAG-FLEX-TdTom at E15.5 to label neurons derived from Tα1-positive and Glast-positive lineages. (**b**) Tissue was collected at P28 where IUE labelled neurons were observed in L2/3 of S1. (**c**) The labelled L2/3 neurons comprised 42.65 % derived from a Tα1-positive/Glast-negative lineage, 29.88 % derived from a Tα1-positive/Glast-positive lineage and 27.48 % derived from a Tα1-negative/Glast-positive lineage (*n* = 8 animals). (**d**) On average, neurons derived from a Tα1-positive lineage resided at 263.34 µm from pia and neurons derived from a Tα1-negative/Glast-positive lineage resided at 244.34 µm from pia (*n* = 8 animals).

**Supplementary Figure 6:**
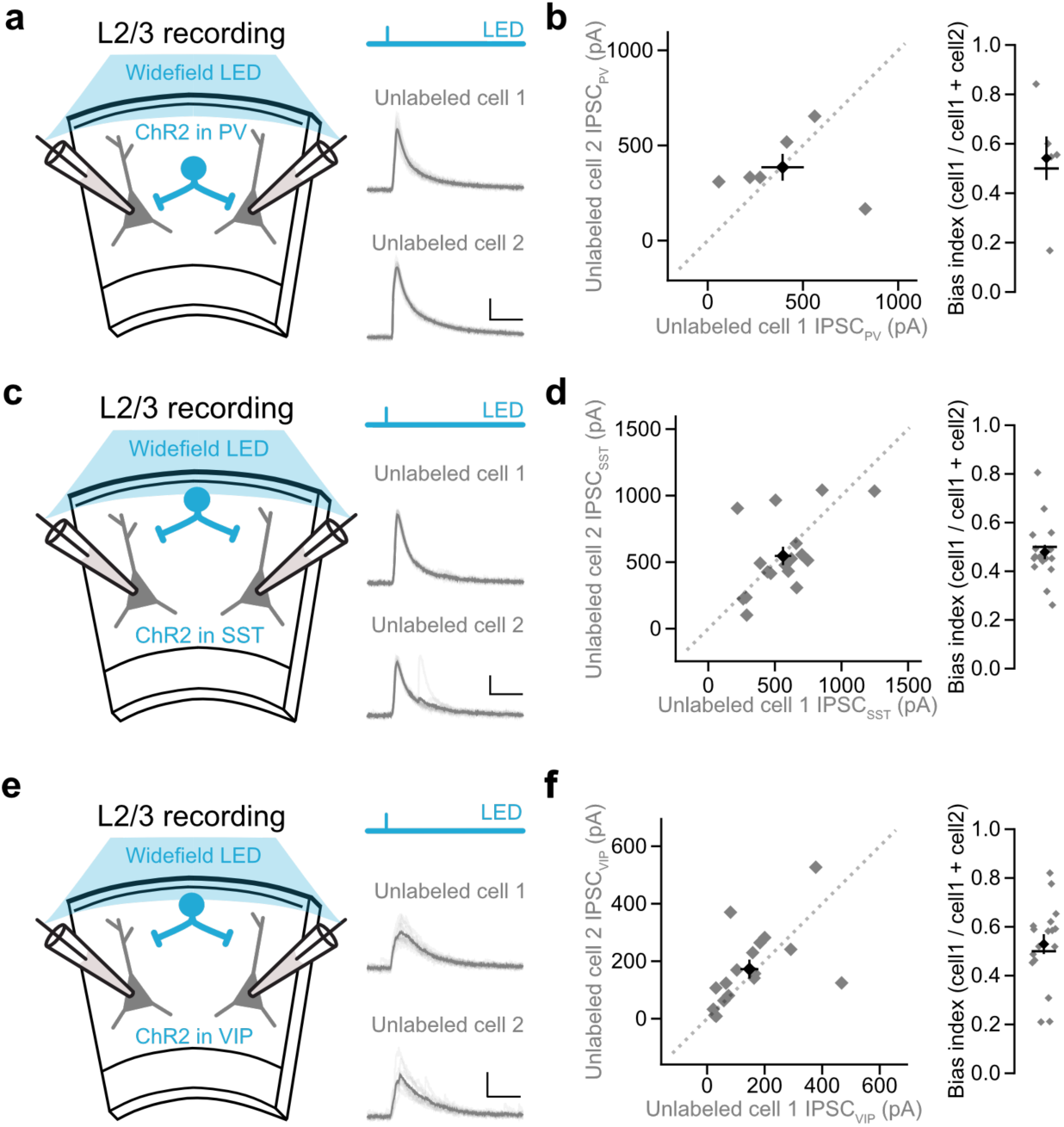
Randomly sampled, unlabeled pairs of L2/3 pyramidal neurons receive similar input from PV, SST and VIP interneurons. (**a**) Acute slices were prepared from S1 and simultaneous recordings were performed from pairs of randomly sampled, unlabeled L2/3 pyramidal neurons. Brief pulses of light from a LED (473 nm, 1ms duration) were used to activate ChR2-expressing PV interneurons throughout the brain slice (left). Example IPSCPV recorded from two unlabeled pyramidal neurons (right). Scale bar 200 pA, 50 ms. (**b**) IPSCPV did not differ between pairs of unlabeled pyramidal neurons (One-sample t-test, unlabeled neuron 1 mean: 393.09 ± 111.52 pA, unlabeled neuron 2 mean: 385.32 ± 70.41 pA, bias index: 0.46 ± 0.09, *n* = 6 pairs from 3 animals, *P* = 0.66, df = 5, T = - 0.47). (**c**) Experimental set up as described in ‘A’, but brief light pulses were used to activate ChR2-expressing SST interneurons throughout the brain slice (left). Example IPSCSST recorded from two unlabeled pyramidal neurons (right). Scale bar 200 pA, 50 ms. (**d**) IPSCSST did not differ between pairs of unlabeled pyramidal neurons (two-tailed Wilcoxon signed-rank test, unlabeled neuron 1 mean: 560.99 ± 62.15 pA, unlabeled neuron 2 mean: 546.70 ± 69.28 pA, bias index = 0.52 ± 0.03, *n* = 17 pairs from 8 animals, *P* = 0.24, W = 51). (**e**) Experimental set up as described in ‘A’, but brief light pulses were used to activate ChR2-expressing VIP interneurons throughout the brain slice (left). Example IPSCVIP recorded from two unlabeled pyramidal neurons (right). Scale bar 100 pA, 50 ms. (**f**) IPSCVIP did not differ between pairs of unlabeled pyramidal neurons (two-tailed one-sample t-test, unlabeled neuron 1 mean: 147.67 ± 27.78 pA, unlabeled neuron 2 mean: 171.96 ± 35.95 pA, bias index = 0.45 ± 0.04, *n* = 17 pairs from 10 animals, *P* = 0.87, df = 16, T = -1.38).

**Supplementary Figure 7:**
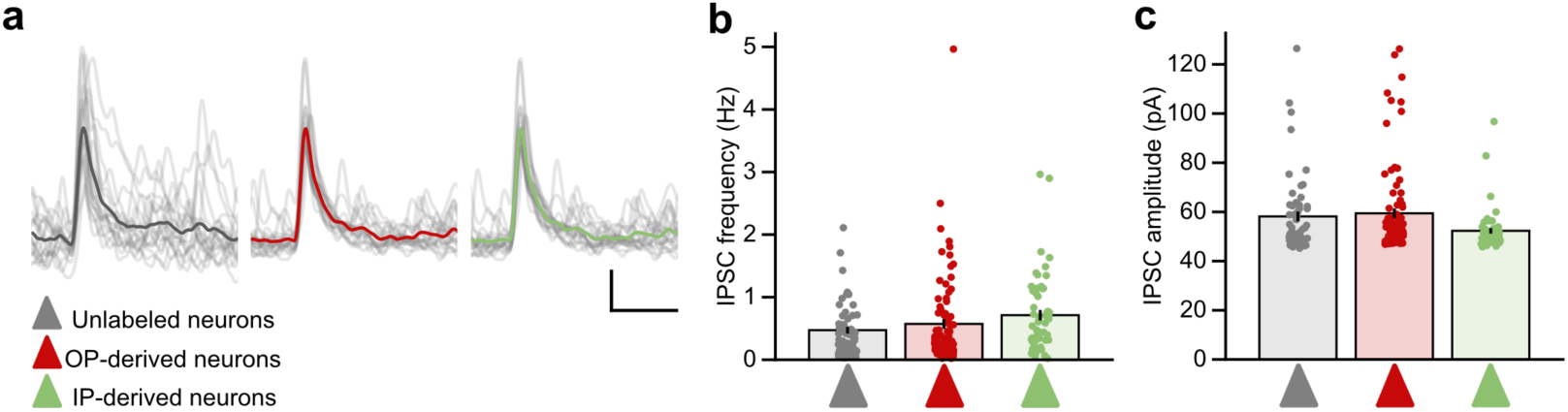
L2/3 pyramidal neurons from different progenitor types receive similar levels of spontaneous action-potential-evoked synaptic inhibition. (**a**) Whole-cell patch clamp recordings were made from unlabeled, OP-derived and IP-derived neurons in cortical slices. IPSCs were detected using automated detection. Example traces show 20 detected IPSCs from each neuron type, mean is overlaid in bold. Scale bar 20 pA, 50 ms. (**b**) IPSC frequency did not differ between unlabeled, OP-derived and IP-derived L2/3 pyramidal neurons (Kruskal-Wallis one-way analysis of variance, unlabeled mean: 0.50 ± 0.06 Hz, *n* = 54 neurons from 16 animals, OP-derived mean: 0.57 ± 0.08 Hz, *n* = 85 neurons from 29 animals, IP-derived mean: 0.71 ± 0.08 Hz, *n* = 55 neurons from 30 animals, *P* = 0.05, H = 5.99). (**c**) IPSC amplitude did not differ between unlabeled, OP-derived and IP-derived L2/3 neurons (Kruskal-Wallis one-way analysis of variance, unlabeled mean: 58.56 ± 2.14 pA, *n* = 54 neurons from 16 animals, OP-derived mean: 59.43 ± 1.99 pA, *n* = 85 neurons from 29 animals, IP-derived mean: 52.31 ± 1.12 pA, *n* = 55 neurons from 30 animals, *P* = 0.08, H = 4.85).

**Supplementary Figure 8:**
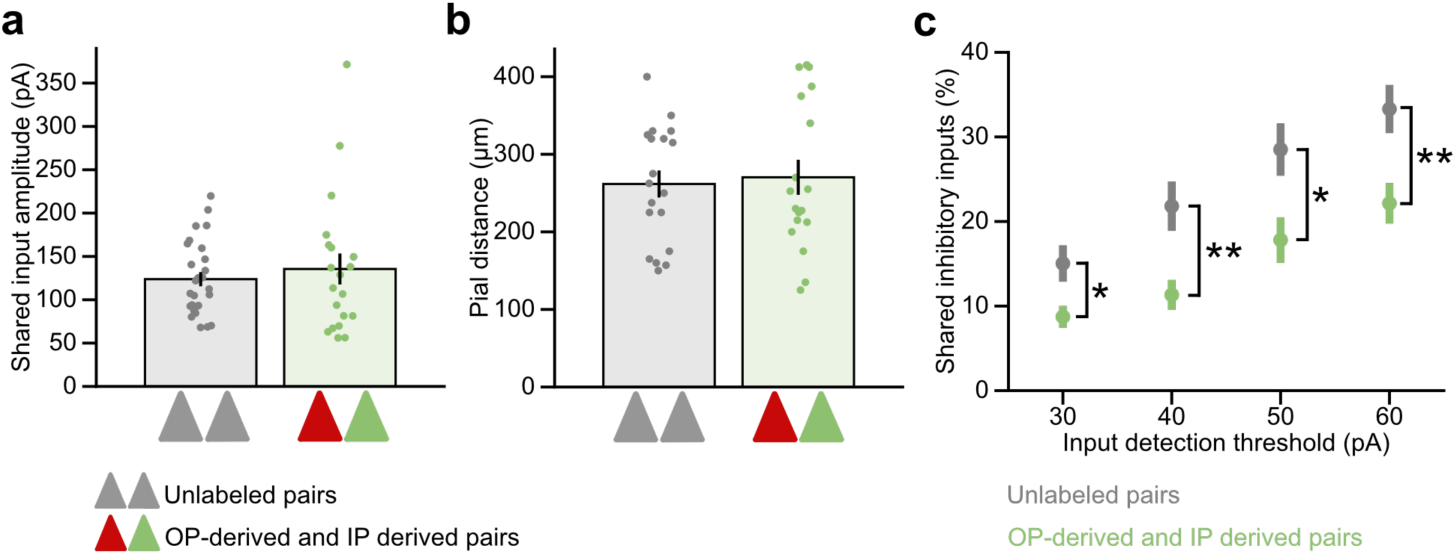
Shared inhibition from spontaneous IPSC recordings does not depend on shared input amplitude, pial distance or input detection threshold. (**a**) Mean amplitude of shared inhibition did not differ between unlabeled pairs and OP-derived and IP-derived pairs (two-tailed Mann-Whitney U test, unlabeled mean: 123.79 ± 8.24 pA, *n* = 27 pairs from 16 animals, OP-derived and IP derived mean: 135.54 ± 17.91 pA, *n* = 20 from 10 animals, *P* = 0.91, U = 276). (**b**) Distance from pia did not differ between unlabeled pairs and OP-derived and IP-derived pairs (two-tailed Mann-Whitney U test, unlabeled mean: 261.68 ± 17.45 µm, *n* = 19 pairs from 13 animals, OP-derived and IP-derived mean: 270.28 ± 17.56 µm, *n* = 18 pairs from 11 animals, *P* = 0.48, U = 164.5). (**c**) OP-derived and IP-derived neuron pairs had less shared inputs than unlabeled pairs across multiple input detection thresholds, ranging from 30 pA to 60 pA. (two-tailed Mann-Whitney U test, 30 pA threshold: *P* = 0.02, U = 157; 40 pA threshold: *P* = 0.007, U = 141; 50 pA threshold: *P* = 0.01, U = 148; 60 pA threshold: *P* = 0.004, U = 133).

**Supplementary Figure 9:**
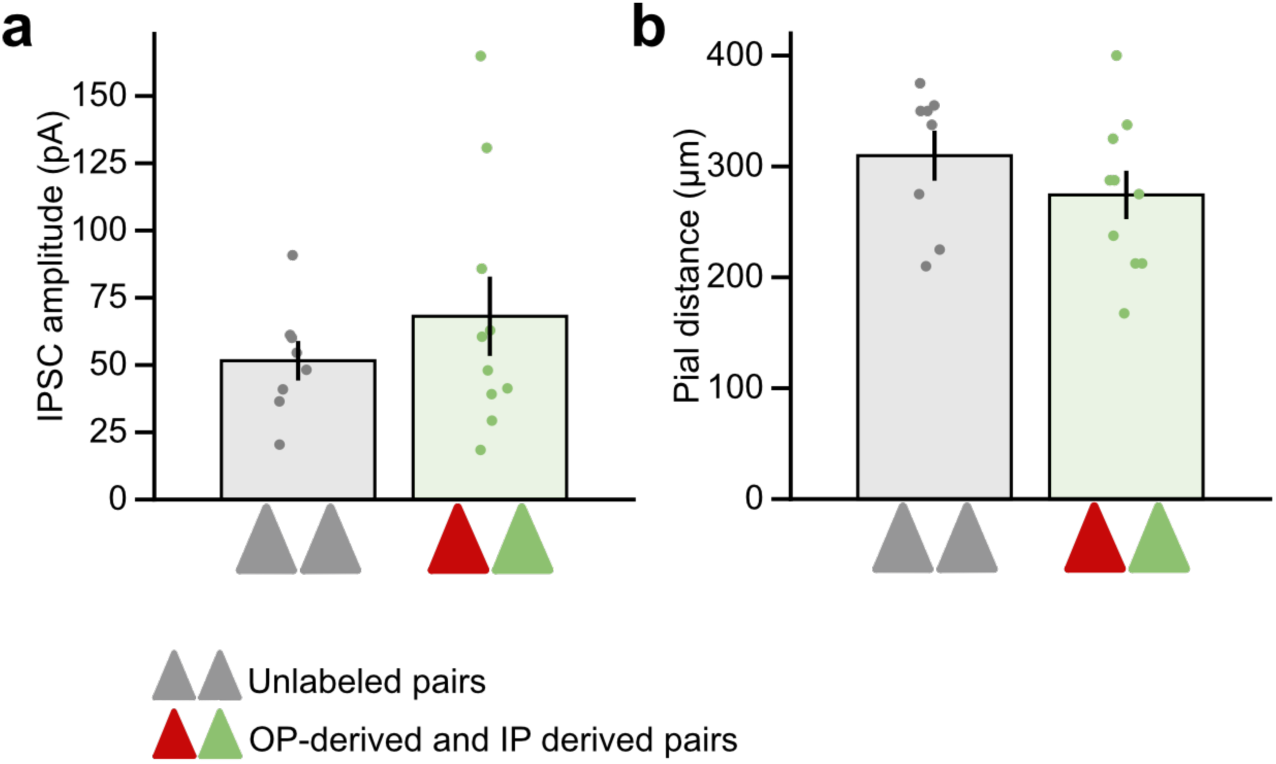
Under minimal optical stimulation, unlabeled pairs and OP-derived and IP-derived pairs do not differ in their IPSC amplitude or pial distance. (**a**) Mean IPSC amplitude does not differ between unlabeled pairs and OP-derived and IP-derived pairs (two-tailed unpaired t-test, unlabeled mean: 51.59 ± 7.36 pA, *n* = 8 pairs from 5 animals, OP-derived and IP-derived mean: 68.10 ± 14.76 pA, *n* = 10 pairs from 6 animals, *P* = 0.37, df = 16, T = -0.93). (**b**) Pial distance does not differ between unlabeled pairs and OP-derived and IP-derived pairs (two-tailed unpaired t-test, unlabeled mean: 309.69 ± 22.62 µm, *n* = 8 pairs from 5 animals, OP-derived and IP-derived mean: 274.25 ± 21.89 µm, *n* = 10 pairs from 6 animals, *P* = 0.28, df = 16, T = 1.12).

## References

1 Pouille, F. & Scanziani, M. Enforcement of temporal fidelity in pyramidal cells by somatic feed-forward inhibition. Science (1979) 293, 1159–1163 (2001).

2. Galarreta, M. & Hestrin, S. Spike transmission and synchrony detection in networks of GABAergic interneurons. Science (1979) 292, 2295–2299 (2001).

3. Panzeri, S., Moroni, M., Safaai, H. & Harvey, C. D. The structures and functions of correlations in neural population codes. Nat Rev Neurosci 23, 551–567 (2022).

4. Williams, L. E. & Holtmaat, A. Higher-Order Thalamocortical Inputs Gate Synaptic Long-Term Potentiation via Disinhibition. Neuron 101, 91–102.e4 (2019).

5. Isaacson, J. S. & Scanziani, M. How inhibition shapes cortical activity. Neuron 72, 231– 243 (2011).

6. Fino, E. & Yuste, R. Dense inhibitory connectivity in neocortex. Neuron 69, 1188–1203 (2011).

7. Packer, A. M. & Yuste, R. Dense, Unspecific Connectivity of Neocortical Parvalbumin-Positive Interneurons: A Canonical Microcircuit for Inhibition? Journal of Neuroscience 31, 13260–13271 (2011).

8. Lee, A. T. et al. Pyramidal neurons in prefrontal cortex receive subtype-specific forms of excitation and inhibition. Neuron 81, 61–68 (2014).

9. Lee, S. H. et al. Parvalbumin-Positive Basket Cells Differentiate among Hippocampal Pyramidal Cells. Neuron 82, 1129–1144 (2014).

10. Varga, C., Lee, S. Y. & Soltesz, I. Target-selective GABAergic control of entorhinal cortex output. Nat Neurosci 13, 822–824 (2010).

11. Krook-Magnuson, E., Varga, C., Lee, S. H. & Soltesz, I. New dimensions of interneuronal specialization unmasked by principal cell heterogeneity. Trends Neurosci 35, 175 (2012).

12. Wu, S. J. et al. Cortical somatostatin interneuron subtypes form cell-type-specific circuits. Neuron 111, 2675–2692.e9 (2023).

13. Jézéquel, J. et al. Cadherins orchestrate specific patterns of perisomatic inhibition onto distinct pyramidal cell populations. bioRxiv 2023.09.28.559922 (2023) doi:10.1101/2023.09.28.559922.

14. Znamenskiy, P. et al. Functional specificity of recurrent inhibition in visual cortex. Neuron 112, 991–1000.e8 (2024).

15. Runyan, C. A. et al. Response features of parvalbumin-expressing interneurons suggest precise roles for subtypes of inhibition in visual cortex. Neuron 67, 847–857 (2010).

16. Kuan, A. T. et al. Synaptic wiring motifs in posterior parietal cortex support decision-making. bioRxiv 2022.04.13.488176 (2022) doi:10.1101/2022.04.13.488176.

17. Jabaudon, D. Fate and freedom in developing neocortical circuits. Nat Commun 8, 1–9 (2017).

18. Greig, L. C., Woodworth, M. B., Galazo, M. J., Padmanabhan, H. & Macklis, J. D. Molecular logic of neocortical projection neuron specification, development and diversity. Nat Rev Neurosci 14, 755–769 (2013).

19. Telley, L. et al. Sequential transcriptional waves direct the differentiation of newborn neurons in the mouse neocortex. Science (1979) 351, 1443–1446 (2016).

20. Telley, L. et al. Temporal patterning of apical progenitors and their daughter neurons in the developing neocortex. Science (1979) 364, (2019).

21. Molyneaux, B. J., Arlotta, P., Menezes, J. R. L. & Macklis, J. D. Neuronal subtype specification in the cerebral cortex. Nat Rev Neurosci 8, 427–437 (2007).

22. Huilgol, D., Russ, J. B., Srivas, S. & Huang, Z. J. The progenitor basis of cortical projection neuron diversity. Curr Opin Neurobiol 81, 102726 (2023).

23. Rakic, P. Specification of Cerebral Cortical Areas. Science (1979) 241, 170–176 (1988).

24. Noctor, S. C., Martinez-Cerdeño, V., Ivic, L. & Kriegstein, A. R. Cortical neurons arise in symmetric and asymmetric division zones and migrate through specific phases. Nat Neurosci 7, 136–144 (2004).

25. Tan, X. & Shi, S. H. Neocortical neurogenesis and neuronal migration. Wiley Interdiscip Rev Dev Biol 2, 443–459 (2013).

26. Gao, P. et al. Deterministic Progenitor Behavior and Unitary Production of Neurons in the Neocortex. Cell 159, 775 (2014).

27. Hippenmeyer, S. Principles of neural stem cell lineage progression: Insights from developing cerebral cortex. Curr Opin Neurobiol 79, 102695 (2023).

28. Noctor, S. C., Flint, A. C., Weissman, T. A., Dammerman, R. S. & Kriegstein, A. R. Neurons derived from radial glial cells establish radial units in neocortex. Nature 409, 714–720 (2001).

29. Noctor, S. C., Martinez-Cerdeño, V., Ivic, L. & Kriegstein, A. R. Cortical neurons arise in symmetric and asymmetric division zones and migrate through specific phases. Nat Neurosci 7, 136–144 (2004).

30. Gal, J. S. et al. Molecular and morphological heterogeneity of neural precursors in the mouse neocortical proliferative zones. Journal of Neuroscience 26, 1045–1056 (2006).

31. Stancik, E. K., Navarro-Quiroga, I., Sellke, R. & Haydar, T. F. Heterogeneity in Ventricular Zone Neural Precursors Contributes to Neuronal Fate Diversity in the Postnatal Neocortex. Journal of Neuroscience 30, 7028–7036 (2010).

32. Kowalczyk, T. et al. Intermediate neuronal progenitors (basal progenitors) produce pyramidal-projection neurons for all layers of cerebral cortex. Cereb Cortex 19, 2439– 2450 (2009).

33. Chenn, A. & McConnell, S. K. Cleavage orientation and the asymmetric inheritance of Notch1 immunoreactivity in mammalian neurogenesis. Cell 82, 631–641 (1995).

34. Ellender, T. J. et al. Embryonic progenitor pools generate diversity in fine-scale excitatory cortical subnetworks. Nat Commun 10, 1–16 (2019).

35. Guillamon-Vivancos, T. et al. Distinct Neocortical Progenitor Lineages Fine-tune Neuronal Diversity in a Layer-specific Manner. Cerebral Cortex 29, 1121 (2019).

36. Tyler, W. A., Medalla, M., Guillamon-Vivancos, T., Luebke, J. I. & Haydar, T. F. Neural precursor lineages specify distinct neocortical pyramidal neuron types. Journal of Neuroscience 35, 6142–6152 (2015).

37. Huilgol, D. et al. Direct and indirect neurogenesis generate a mosaic of distinct glutamatergic projection neuron types in cerebral cortex. Neuron 111, (2023).

38. Buchan, M. J. et al. Higher-order thalamocortical circuits are specified by embryonic cortical progenitor types in the mouse brain. Cell Rep 43, (2024).

39. Rakic, P. Specification of Cerebral Cortical Areas. Science (1979) 241, 170–176 (1988).

40. Wichterle, H., Turnbull, D. H., Nery, S., Fishell, G. & Alvarez-Buylla, A. In utero fate mapping reveals distinct migratory pathways and fates of neurons born in the mammalian basal forebrain. Development 128, 3759–3771 (2001).

41. Nery, S., Fishell, G. & Corbin, J. G. The caudal ganglionic eminence is a source of distinct cortical and subcortical cell populations. Nat Neurosci 5, 1279–1287 (2002).

42. Anderson, S. A., Kaznowski, C. E., Horn, C., Rubenstein, J. L. R. & McConnell, S. K. Distinct origins of neocortical projection neurons and interneurons in vivo. Cerebral Cortex 12, 702–709 (2002).

43. Anthony, T. E., Klein, C., Fishell, G. & Heintz, N. Radial glia serve as neuronal progenitors in all regions of the central nervous system. Neuron 41, 881–890 (2004).

44. Gal, J. S. et al. Molecular and Morphological Heterogeneity of Neural Precursors in the Mouse Neocortical Proliferative Zones. Journal of Neuroscience 26, 1045–1056 (2006).

45. Stancik, E. K., Navarro-Quiroga, I., Sellke, R. & Haydar, T. F. Heterogeneity in Ventricular Zone Neural Precursors Contributes to Neuronal Fate Diversity in the Postnatal Neocortex. Journal of Neuroscience 30, 7028–7036 (2010).

46. Kowalczyk, T. et al. Intermediate Neuronal Progenitors (Basal Progenitors) Produce Pyramidal–Projection Neurons for All Layers of Cerebral Cortex. Cerebral Cortex 19, 2439–2450 (2009).

47. Chenn, A. & McConnell, S. K. Cleavage orientation and the asymmetric inheritance of Notch1 immunoreactivity in mammalian neurogenesis. Cell 82, 631–641 (1995).

48. Langevin, L. M. et al. Validating in utero electroporation for the rapid analysis of gene regulatory elements in the murine telencephalon. Developmental Dynamics 236, 1273–1286 (2007).

49. Gal, J. S. et al. Molecular and Morphological Heterogeneity of Neural Precursors in the Mouse Neocortical Proliferative Zones. Journal of Neuroscience 26, 1045–1056 (2006).

50. Mizutani, K. I., Yoon, K., Dang, L., Tokunaga, A. & Gaiano, N. Differential Notch signalling distinguishes neural stem cells from intermediate progenitors. Nature 449, 351–355 (2007).

51. Stancik, E. K., Navarro-Quiroga, I., Sellke, R. & Haydar, T. F. Heterogeneity in ventricular zone neural precursors contributes to neuronal fate diversity in the postnatal neocortex. Journal of Neuroscience 30, 7028–7036 (2010).

52. Haider, B., Schulz, D. P. P. A., Häusser, M. & Carandini, M. Millisecond Coupling of Local Field Potentials to Synaptic Currents in the Awake Visual Cortex. Neuron 90, 35–42 (2016).

53. Okun, M. et al. Diverse coupling of neurons to populations in sensory cortex. Nature 521, 511–515 (2015).

54. Okun, M., Naim, A. & Lampl, I. The Subthreshold Relation between Cortical Local Field Potential and Neuronal Firing Unveiled by Intracellular Recordings in Awake Rats. Journal of Neuroscience 30, 4440–4448 (2010).

55. Cutts, C. S. & Eglen, X. S. J. Detecting pairwise correlations in spike trains: an objective comparison of methods and application to the study of retinal waves. The Journal of Neuroscience 34, 14288–14303 (2014).

56. Ellender, T. J. et al. Embryonic progenitor pools generate diversity in fine-scale excitatory cortical subnetworks. Nat Commun 10, 1–16 (2019).

57. Markram, H. et al. Interneurons of the neocortical inhibitory system. Nat Rev Neurosci 5, 793–807 (2004).

58. Fino, E., Packer, A. M. & Yuste, R. The logic of inhibitory connectivity in the neocortex. Neuroscientist vol. 19 228–237 Preprint at 10.1177/1073858412456743 (2013).

59. Xue, M., Atallah, B. V. & Scanziani, M. Equalizing excitation-inhibition ratios across visual cortical neurons. Nature 511, 596–600 (2014).

60. Markram, H. et al. Interneurons of the neocortical inhibitory system. Nature Reviews Neuroscience vol. 5 793–807 Preprint at 10.1038/nrn1519 (2004).

61. Pfeffer, C. K., Xue, M., He, M., Huang, Z. J. & Scanziani, M. Inhibition of inhibition in visual cortex: the logic of connections between molecularly distinct interneurons. Nat Neurosci 16, 1068–1076 (2013).

62. Naskar, S., Qi, J., Pereira, F., Gerfen, C. R. & Lee, S. Cell-type-specific recruitment of GABAergic interneurons in the primary somatosensory cortex by long-range inputs. Cell Rep 34, 108774 (2021).

63. Sippy, T. & Yuste, R. Decorrelating action of inhibition in neocortical networks. The Journal of Neuroscience 33, 9813–9830 (2013).

64. Safari, M. S., Mirnajafi-Zadeh, J., Hioki, H. & Tsumoto, T. Parvalbumin-expressing interneurons can act solo while somatostatin-expressing interneurons act in chorus in most cases on cortical pyramidal cells. Sci Rep 7, 1–14 (2017).

65. Morgenstern, N. A., Bourg, J. & Petreanu, L. Multilaminar networks of cortical neurons integrate common inputs from sensory thalamus. Nat Neurosci 19, 1034–1040 (2016).

66. Hasenstaub, A. et al. Inhibitory postsynaptic potentials carry synchronized frequency information in active cortical networks. Neuron 47, 423–435 (2005).

67. Haider, B., Schulz, D. P. P. A., Häusser, M. & Carandini, M. Millisecond Coupling of Local Field Potentials to Synaptic Currents in the Awake Visual Cortex. Neuron 90, 35–42 (2016).

68. Atallah, B. V. & Scanziani, M. Instantaneous Modulation of Gamma Oscillation Frequency by Balancing Excitation with Inhibition. Neuron 62, 566–577 (2009).

69. Ruan, X. et al. Progenitor cell diversity in the developing mouse neocortex. Proc Natl Acad Sci U S A 118, e2018866118 (2021).

70. Li, Z. et al. Transcriptional priming as a conserved mechanism of lineage diversification in the developing mouse and human neocortex. Sci Adv 6, (2020).

71. Tasic, B. et al. Shared and distinct transcriptomic cell types across neocortical areas. Nature 563, 72–78 (2018).

72. Di Bella, D. J. et al. Molecular logic of cellular diversification in the mouse cerebral cortex. Nature 595, 554–559 (2021).

73. Tamamaki, N., Nakamura, K., Okamoto, K. & Kaneko, T. Radial glia is a progenitor of neocortical neurons in the developing cerebral cortex. Neurosci Res 41, 51–60 (2001).

74. Noctor, S. C., Martinez-Cerdeño, V., Ivic, L. & Kriegstein, A. R. Cortical neurons arise in symmetric and asymmetric division zones and migrate through specific phases. Nat Neurosci 7, 136–144 (2004).

75. Martínez-Cerdeño, V., Noctor, S. C. & Kriegstein, A. R. The role of intermediate progenitor cells in the evolutionary expansion of the cerebral cortex. Cerebral Cortex 16 **Suppl 1**, (2006).

76. Florio, M. & Huttner, W. B. Neural progenitors, neurogenesis and the evolution of the neocortex. Development 141, 2182–2194 (2014).

77. Matho, K. S. et al. Genetic dissection of the glutamatergic neuron system in cerebral cortex. Nature 2021 598:7879 598, 182–187 (2021).

78. Matho, K. S. et al. Genetic dissection of the glutamatergic neuron system in cerebral cortex. Nature 598, 182–187 (2021).

79. Tasic, B. et al. Shared and distinct transcriptomic cell types across neocortical areas. Nature 563, 72–78 (2018).

80. Gouwens, N. W. et al. Integrated Morphoelectric and Transcriptomic Classification of Cortical GABAergic Cells. Cell 183, 935–953.e19 (2020).

81. Helmstaedter, M., Sakmann, B. & Feldmeyer, D. Neuronal correlates of local, lateral, and translaminar inhibition with reference to cortical columns. Cerebral Cortex 19, 926–937 (2009).

82. Kätzel, D., Zemelman, B. V., Buetfering, C., Wölfel, M. & Miesenböck, G. The columnar and laminar organization of inhibitory connections to neocortical excitatory cells. Nat Neurosci 14, 100–107 (2010).

83. Yoshimura, Y., Dantzker, J. L. M. & Callaway, E. M. Excitatory cortical neurons form fine-scale functional networks. Nature 433, 868–873 (2005).

84. Denardo, L. A., Berns, D. S., Deloach, K. & Luo, L. Connectivity of mouse somatosensory and prefrontal cortex examined with trans-synaptic tracing. Nat Neurosci 18, 1687– 1697 (2015).

85. Huszár, R., Zhang, Y., Blockus, H. & Buzsáki, G. Preconfigured dynamics in the hippocampus are guided by embryonic birthdate and rate of neurogenesis. Nat Neurosci 25, 1201–1212 (2022).

86. Cavalieri, D. et al. Ca1 pyramidal cell diversity is rooted in the time of neurogenesis. Elife 10, (2021).

87. Xu, H. T. et al. Distinct lineage-dependent structural and functional organization of the hippocampus. Cell 157, 1552 (2014).

88. Harris, K. D. & Shepherd, G. M. G. The neocortical circuit: themes and variations. Nat Neurosci 18, 170 (2015).

89. Mercier, B. E., Legg, C. R. & Glickstein, M. Basal ganglia and cerebellum receive different somatosensory information in rats. Proc Natl Acad Sci U S A 87, 4388–4392 (1990).

90. Huilgol, D. et al. Direct and indirect neurogenesis generate a mosaic of distinct glutamatergic projection neuron types in cerebral cortex. Neuron 111, 2557–2569.e4 (2023).

91. Tyler, W. A. & Haydar, T. F. Multiplex genetic fate mapping reveals a novel route of neocortical neurogenesis, which is altered in the Ts65Dn mouse model of Down syndrome. The Journal of Neuroscience 33, 5106–5119 (2013).

92. Margrie, T. W., Brecht, M. & Sakmann, B. In vivo, low-resistance, whole-cell recordings from neurons in the anaesthetized and awake mammalian brain. Pflugers Arch 444, 491–498 (2002).

93. Wang, Y., Liu, Y. Z., Wang, S. Y. & Wang, Z. In vivo whole-cell recording with high success rate in anaesthetized and awake mammalian brains. Mol Brain 9, 1–14 (2016).

94. Jordan, R. Optimized protocol for in vivo whole-cell recordings in head-fixed, awake behaving mice. STAR Protoc 2, (2021).

95. Traynelis, S. F. Software-based correction of single compartment series resistance errors. J Neurosci Methods 86, 25–34 (1998).

96. Cutts, C. S. & Eglen, X. S. J. Detecting pairwise correlations in spike trains: an objective comparison of methods and application to the study of retinal waves. The Journal of Neuroscience 34, 14288–14303 (2014).

97. Chini, M., Pfeffer, T. & Hanganu-Opatz, I. An increase of inhibition drives the developmental decorrelation of neural activity. Elife 11, (2022).

98. Yang, W. et al. Anesthetics fragment hippocampal network activity, alter spine dynamics, and affect memory consolidation. PLoS Biol 19, e3001146 (2021).

99. Yoshimura, Y. & Callaway, E. M. Fine-scale specificity of cortical networks depends on inhibitory cell type and connectivity. Nature Neuroscience 2005 8:11 8, 1552–1559 (2005).

100. Morgenstern, N. A., Bourg, J. & Petreanu, L. Multilaminar networks of cortical neurons integrate common inputs from sensory thalamus. Nature Neuroscience 2016 19:8 19, 1034–1040 (2016).

